# A Kinetochore-Associated Proteasome Pool Drives a Second Pathway of Cohesin Removal during Meiosis

**DOI:** 10.64898/2026.08.18.745540

**Authors:** Anshul Mishra, N. Rory Butler, Lori B. Koch, Christos Spanos, Aaron F. Severson, Adele L. Marston, G. Valentin Börner

## Abstract

Accurate chromosome segregation requires the spatiotemporally regulated removal of sister chromatid cohesion. Cohesin cleavage by the endopeptidase separase depends on destruction of its inhibitor securin by the ubiquitin-proteasome system (UPS) and on phosphorylation-mediated priming of the cohesin kleisin subunit. Whether the UPS also contributes to cohesin priming has remained unknown. Here, we show that the 26*S* proteasome mediates cohesin removal during meiosis II through branches of two parallel pathways. Using separation-of-function mutants targeting either the proteasome’s core or regulatory particles, we identify a proteasome function that is required specifically for centromeric cohesin removal during meiosis II but dispensable for separase activation. Defects in this proteasome function causes centromeric accumulation of phosphatase anchor shugoshin (Sgo1), impaired cleavage of meiotic kleisin Rec8, and frequent failure of sister chromatid segregation. Bypassing the requirement for Rec8 priming, either through a phosphomimetic *rec8* allele or by separase-independent Rec8 cleavage, restores chromosome segregation, demonstrating that the proteasome mediates cohesin removal independently of its established role in activating separase. Consistent with a direct role, proteasomes localize prominently to kinetochores during meiosis II. Together, these findings identify the proteasome as a dual-function regulator that mediates both separase activation and cohesin priming, revealing how a single proteolytic machine coordinates the two molecular pathways underlying stepwise chromosome segregation during meiosis.

## Introduction

The ubiquitin-proteasome system is a major regulator of eukaryotic cell division. By selectively degrading key cell-cycle proteins, the 26*S* proteasome coordinates replication, recombination, and chromosome segregation ^1^. One of the UPS’s fundamental functions is the control of chromosome segregation through activation of the cohesin-cleaving endopeptidase separase ^2^. At anaphase onset, the anaphase-promoting complex/cyclosome (APC/C) polyubiquitinates the separase inhibitor securin, marking it for degradation and thereby activating separase to cleave cohesin and trigger chromosome disjunction ^3^. While this pathway is well established, it is unknown whether the proteasome contributes to cohesin removal in additional ways.

The 26*S* proteasome is a multisubunit protease that degrades proteins earmarked for destruction within its enclosed proteolytic chamber ^4^. It comprises the barrel-shaped core particle (CP, 20*S*), which is associated with one or two regulatory particles (RP, 19*S*). The CP consists of two stacked, heptameric β-rings sandwiched between two heptameric α-rings. The β-rings harbor three endopeptidase activities ^5^, while the α-rings regulate client protein access to the proteolytic chamber ^6^. The RP docks onto, deubiquitinates, and unfolds client proteins to feed them into the CP ^4^. Client protein unfolding is driven by a ring of six ATPases (Rpt1 to 6) at the base of the RP ^7,8^.

The cohesin complex holds the two DNA duplexes together after replication, conferring the sister chromatid cohesion required for timely and accurate chromosome segregation by counteracting the pulling forces of the spindle. The spatial and temporal regulation of cohesin removal determines the mode of chromosome segregation in both mitosis and meiosis ^9^. In mitosis, sister chromatid segregation involves cohesin removal along the length of chromosomes, achieved in a single round in yeast ^2^ and in two steps in vertebrates ^10,11^. In meiosis, cohesin is cleaved by separase in two sequential steps ^12,13^. During meiosis I, arm cohesion together with crossovers formed by Spo11-induced double-strand breaks, link parental chromosomes (homologues), which segregate upon cleavage of arm cohesin at the metaphase I–anaphase I transition ^14,15^. Pericentromeric cohesin is protected at this stage, maintaining sister chromatid cohesion until its cleavage at the metaphase II–anaphase II transition enables sister chromatid segregation ^16–18^.

The cohesin ring comprises Smc1 and Smc3 as well as a kleisin component that differs between mitotic and meiotic cells ^19^. Whereas mitotic cells utilize kleisin Scc1^Rad21^, meiotic cells incorporate Rec8, whose regulated cleavage underlies the two-step pattern of chromosome segregation characteristic of meiosis ^12,20^. Rec8 cleavage is controlled by two complementary pathways. First, APC/C-dependent degradation of securin activates separase ^3,21^. Second, phosphorylation of Rec8 by several kinases primes it for separase-mediated cleavage ^14,15^.

Protection of centromeric Rec8 during meiosis I is achieved through localized antagonism to priming phosphorylation. Shugoshin (Sgo1 in yeast and Sgo2 in mammals) recruits protein phosphatase 2A (PP2A) to pericentromeric regions, where PP2A removes Rec8 phosphorylation thereby protecting it from cleavage ^22–25^. Additional protection is provided by the meiosis I regulator Spo13^meikin^, which counteracts Rec8 kinases at centromeres^26^. At the meiosis II metaphase-to-anaphase transition, Sgo1 and PP2A are displaced from centromeres, allowing Rec8 phosphorylation by casein kinase 1δ’s catalytic component (Hrr25) and separase cleavage ^16^. Thus, the meiosis I-to-meiosis II transition requires not only separase activity but also the removal of factors that counter priming phosphorylation of centromeric Rec8.

Dissecting roles of the proteasome in chromosome segregation has been challenging because most of its components are essential and broadly required for protein turnover. One exception is the constitutive, though non-essential, α3 subunit of the proteasome CP, encoded by *PRE9* in yeast. In α3^Pre9^’s absence, a second copy of α4^Pre6^ occupies the vacant position, allowing assembly of a functional but altered proteasome ^27,28^. Though dispensable for vegetative growth, α3-containing proteasomes are required specifically during meiotic prophase I where they mediate homologue pairing, synapsis, and recombination ^29,30^. During prophase I, proteasome components further localize to chromosomes in an evolutionarily conserved manner, suggesting that proteasomal activity can be deployed directly at chromosomal sites rather than exclusively through stochastic substrate encounters ^29,30^.

Here, we investigated proteasome roles in mitosis and meiosis using separation-of-function mutants that impair 20*S* core particle and 19*S* regulatory particle functions. We uncover a proteasome function that is specifically required for meiosis II sister chromatid segregation but dispensable for cohesin removal during meiosis I and mitosis. Our findings demonstrate that this novel proteasome role in meiosis II promotes displacement of Sgo1^shugoshin^, deprotection and cleavage of centromeric Rec8, and efficient sister chromatid segregation during meiosis II. These findings reveal that the proteasome controls cohesin removal during meiosis via two parallel pathways.

## Results

### Different proteasome defects delay meiosis I and meiosis II

To track meiotic progression in synchronized budding yeast cultures, we initially distinguished three key stages: mononucleate cells (pre-meiotic S-phase to metaphase I), binucleate cells produced by meiosis I (anaphase I to metaphase II), and tetranucleate cells produced by meiosis II (anaphase II to tetrad formation, when the four nuclei have been encapsulated within spore walls) ^31^. In cells lacking the α3^Pre9^ proteasome subunit, progression through meiosis is delayed at temperatures that do not trigger arrest (e.g., 23°C) ^29^; however, underlying mechanisms and contributions of meiosis I and meiosis II to the delay have not been examined.

To dissect this, we assessed the role of *SPO11*-induced events, including DSB processing and homologue synapsis, on meiotic progression in *pre9Δ*. Cumulative plots of binucleate and tetranucleate cell formation revealed that *pre9Δ* cells exhibit an approximately 2.5-hour delay in completing meiosis I relative to wild type (**Fig. 1A-i**) ^29^. This delay was abolished in a mutant that fails to form DSBs due to an inactivating substitution at Spo11’s catalytic center (*spo11-Y135F-HA_3_His_6_,* hereafter *spo11-yf*; **Fig. 1A-ii**). Thus, the meiosis I delay in *pre9Δ* is triggered by defects in prophase I events initiated by programmed DSBs.

**Fig. 1.**
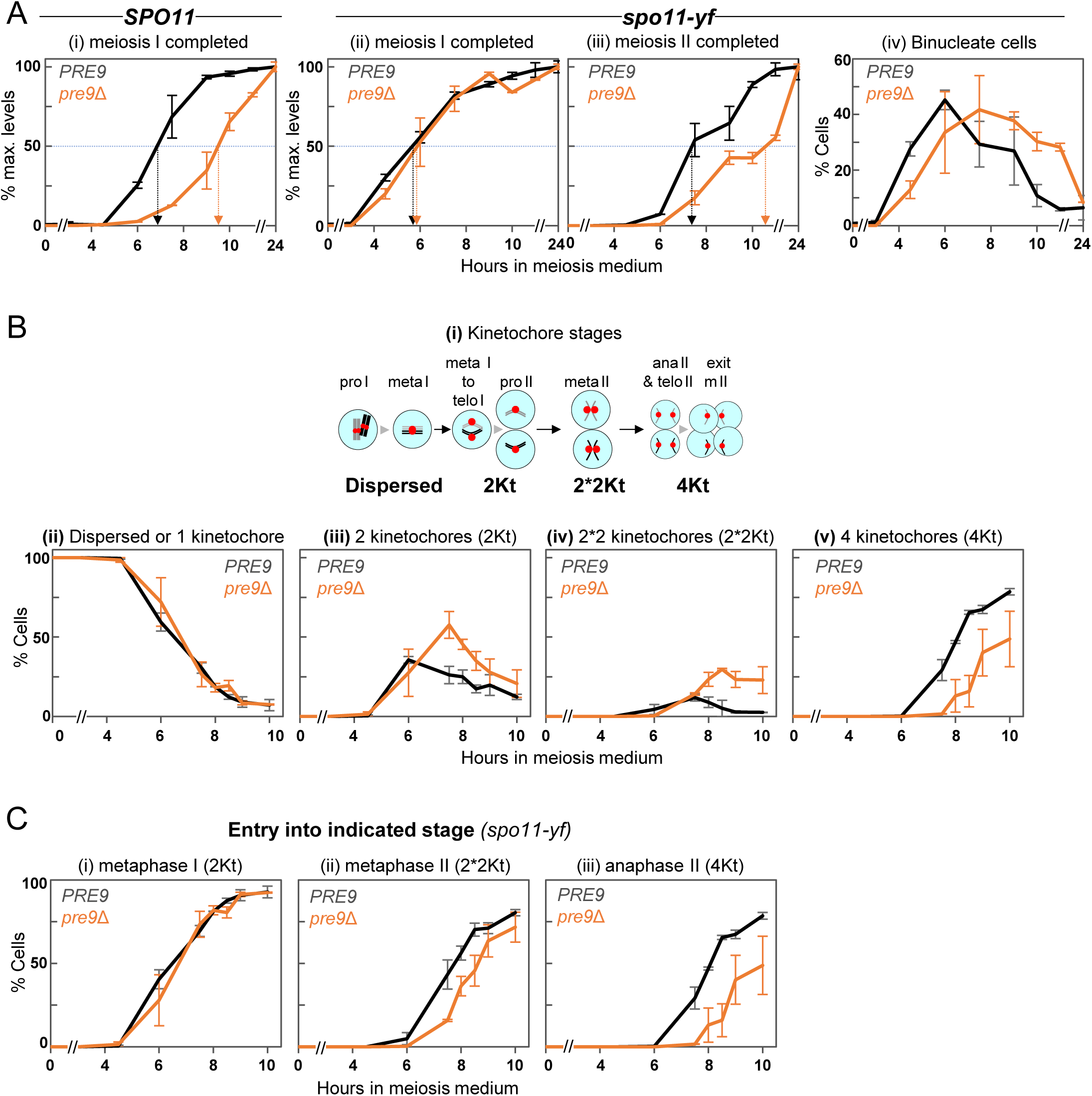
A distinct role of the proteasome in meiosis II progression. (A) The meiosis I delay observed in *pre9Δ* is eliminated in absence of *SPO11-*initiated events, whereas a delay in meiosis II persists. Effects of *pre9Δ* were analyzed **(i)** in *SPO11* cells for the kinetics of meiosis I nuclear divisions, and in *spo11-yf* cells for **(ii)** the kinetics of cumulative meiosis I nuclear divisions, **(iii)** the kinetics of meiosis II nuclear divisions, and **(iv)** the steady state levels of binucleate cells. Completion of meiosis I is defined by the appearance of cells with two or more nuclei, while meiosis II progression is defined by the appearance of cells with more than two nuclei, as assessed by DAPI staining. Graphs (i) to (iii) show values normalized to the maximum percentage, whereas graph (iv) presents steady state levels. Error bars indicate ranges (n = 2). The horizontal dotted line marks 50% of maximum levels, and vertical arrows indicate the time at which 50% of cells have entered the respective stage. Absolute values for each stage are shown in **Fig. S1**. (B) A proteasome requirement specific to meiosis II as revealed by analysis of kinetochore dynamics in wild-type *PRE9* and *pre9Δ* meiotic cultures also carrying *spo11-yf*. **(i)** Schematic depiction of meiotic cell division stages from prophase I to completion of meiosis II. One representative homologue pair in black and grey, kinetochores are in red, nuclei in light blue. **(ii)** Prophase I and early metaphase I: Kinetochore signals dispersed across nucleoplasm or compacted at a single position (1Kt); **(iii)** later metaphase I to prophase II: two compact kinetochore groups at various distances (2Kt); **(iv)** metaphase II: two closely juxtaposed kinetochore pairs, where the two foci are separated by < 1 μm (2*2Kt); **(v)** anaphase II and subsequent stages indicated by four kinetochore groups with no obvious grouping (4Kt). Kinetochores are marked by fluorescently-tagged kinetochore protein Mtw1(-tdTomato). For representative images, see Fig. 4C**-i**. Averages of two parallel cultures are shown, error bars indicate ranges. (C) Cumulative absolute levels of the indicated stages including all later appearing stages for meiotic cultures shown in Fig. 1B.

When focusing on the timing of meiosis II in the same *spo11-yf* background, *pre9Δ* cells still exhibited a ∼3-hour delay, with 50% of cells completing meiosis II at t ∼ 10.5 h in *pre9Δ* compared to ∼7.5 h in wild-type *PRE9* (**Fig. 1A-iii**). Delayed completion of meiosis II was also indicated by a transient accumulation of binucleate cells in *pre9Δ* (**Fig. 1A-iv**). Binucleate cells likely also accumulate transiently in a *SPO11* background, but this was not detected previously due to a sampling gap between 12 h and 24 h ^29^.

Binucleate cells identified by chromatin-staining encompass multiple meiotic stages, ranging from anaphase I to metaphase II. To pinpoint stage-specific delays, we monitored configurations of kinetochores fluorescently tagged with Mtw1-tdTomato (Mtw1-tdT hereafter) in fixed cells. Four cytologically distinguishable stages of meiosis I and II were most abundant in the *spo11-yf* background (see **Fig. 1B-i**). (1) the ‘dispersed’ stage, corresponding to prophase I, when kinetochores are spread throughout the nucleus, and prometaphase I; (2) the ‘2Kt’ stage, corresponding to late metaphase I, anaphase I, telophase I and prophase II, when kinetochore signals have coalesced into two foci spaced one to six μm apart; (3) the ‘2*2Kt’ stage, corresponding to metaphase II, when each kinetochore focus has split into a closely associated pair of foci (less than one μm apart); and (4) the ‘4Kt’ stage, representing anaphase II or later, when the four kinetochore foci are widely separated (greater than one μm apart). To pinpoint the exact meiosis II stage(s) responsible for the delay, progression analysis was performed in the *spo11-yf* background which eliminates the *pre9Δ* meiosis I delay (**Fig. 1A**).

**Figures 1B and 1C** show steady-state levels and cumulative entry curves, respectively, for distinct kinetochore classes. In *pre9Δ*, cells with dispersed or single kinetochores disappeared with kinetics and efficiency indistinguishable from wild-type *PRE9,* consistent with normal exit from prophase I (**Fig. 1B-ii**). Entry into metaphase I also was unaffected, as indicated by appearance with normal timing of cells with two kinetochores (2Kt; **Fig. 1C-i**). However, cells with two kinetochores persisted transiently in *pre9Δ*, resulting in two-fold higher steady-state levels than in wild-type *PRE9*, suggesting delayed progression into metaphase II (**Fig. 1B-iii**). Consistent with this interpretation, entry into metaphase II (2*2Kt) was delayed by one hour in *pre9Δ*, and anaphase II cells (4Kt) appeared with an additional delay of one hour (**Fig. 1C-ii, iii**). Finally, ∼1/3 of *pre9Δ* cells were transiently or permanently arrested in meiosis II at t = 10 h, as indicated by a two-fold increase in steady-state levels of cells blocked at the 2Kt stage (i.e. prior to exit from prophase II), a five-fold increase in cells arrested in metaphase II compared to wild-type *PRE9* (**Fig. 1B-iii, iv**), and an overall decrease in cells that complete meiosis (**Fig. S1**).

Together, these findings indicate that the meiosis I delay in *pre9Δ* is driven by defects in processing of *SPO11*-induced DSBs and associated prophase I events, whereas the meiosis II progression defect arises from a distinct, DSB-independent mechanism. Delayed exit from prophase II and from metaphase II are the major contributors to the *pre9Δ* meiosis II delay*Δ* each accounting for 1 to 1.5 hours.

### The α3^Pre9^ proteasome subunit is required for accurate chromosome segregation exclusively during meiosis II

*pre9Δ* cells produce aneuploid gametes with reduced viability at 23°C, i.e. a temperature that allows tetrad formation ^29^. To determine the cause of aneuploidy in *pre9Δ*, we monitored chromosome segregation during meiosis I, meiosis II, and mitosis. Unless noted, diploid wild-type *SPO11* cultures carried a centromere-linked GFP tag on one or both copies of either chromosome 5 or chromosome 3, with both chromosomes yielding similar results (**Fig. 2**).

**Fig. 2.**
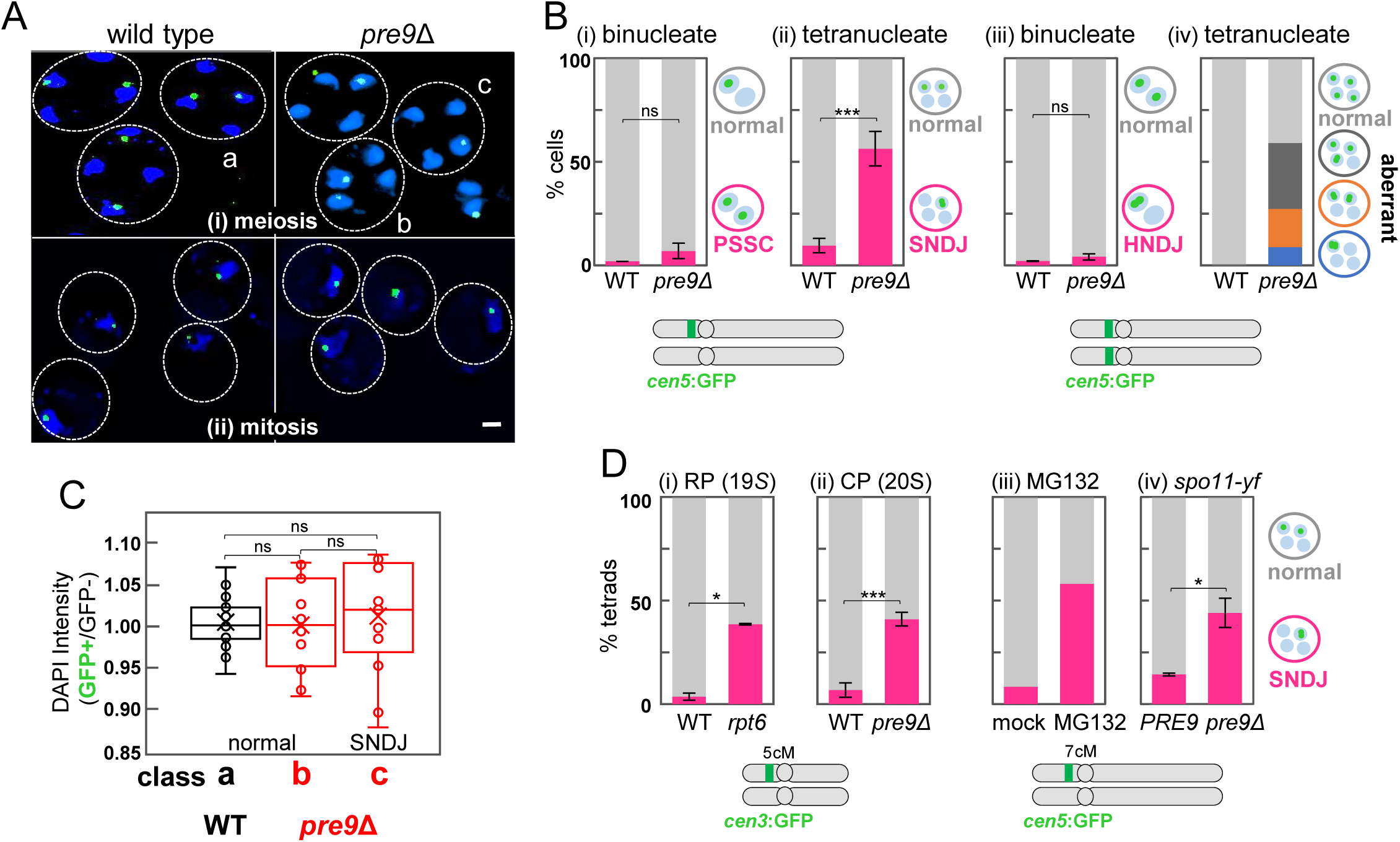
Meiosis II sister chromatid disjunction is distinctly sensitive to impaired proteasome function. (A) Effects of impaired proteasome function on sister chromatid segregation during meiosis and mitosis. Representative fields of yeast cells heterozygous for *cen5*-GFP/*cen5* stained with DAPI in wild type (left) and *pre9Δ* (right). **(i)** Tetranucleate cells after completion of meiosis II cell division. Letters indicate cells exhibiting normal (a, b) or aberrant (c) segregation of GFP-tagged sister chromatids. **(ii)** G1-arrested cells prior to induction of meiosis. Mitotic sister chromatid non-disjunction during premeiotic growth should generate a viable subpopulation lacking a GFP signal which was not observed. Dotted lines indicate cell wall. Size bar is 1 μm. (B) An á3^Pre9^-containing proteasome is specifically required for meiosis II sister chromatid segregation. Bar graphs indicate frequencies of **(i)** precocious separation of sister chromatids (PSSC) during meiosis I, as measured in binucleate cells heterozygous for *cen5*-GFP [WT (N = 2, n = 161); *pre9Δ* (N = 2, n = 121)]; **(ii)** meiosis II sister chromatid nondisjunction (SNDJ) in tetranucleate cells of the same genotypes [WT (N = 4, n = 369); *pre9Δ* (N = 4, n = 287)]. **(iii)** Cells homozygous for *cen5*-GFP were assessed for meiosis I homologue nondisjunction (HNDJ) in binucleate cells [WT (N = 2, n = 149); *pre9Δ* (N = 2, n = 95)], and **(iv)** for overall chromosome 5 missegregation (aberrant) in tetranucleate cells [WT (n = 74); *pre9Δ* (n = 93)]. N and n indicate numbers of meiotic cultures and cells, respectively. Error bars indicate ranges. Schematic depiction of chromosomes 5 with centromeres and TetR-GFP binding sites. (C) Meiosis I SNDJ in cells heterozygous for *cen5*-GFP/*cen5* is not associated with gross chromatin enrichment in GFP-positive nuclei compared to GFP-negative nuclei. Labels a (n = 20), b (n = 13), and c (n = 14) refer to cell classes shown in Fig. 2A**(i)**. Measurements are ratios of average DAPI intensities of nuclei containing GFP versus those lacking GFP in the same cell. Values >1 are expected if chromatin was preferentially segregating into nuclei that have received both copies of *cen5*-GFP. The wider range of ratios in *pre9Δ* could be due to excess of deficit of multiple chromosomes. Box-and-whisker plots show the median (center line), the interquartile range (box), and whiskers extending to 1.5 × interquartile range. Significance was determined by Tukey’s multiple comparison test in one-way anova. (D) Frequency of meiosis II sister chromatid non-disjunction in tetrads derived from cells heterozygous for GFP at *cen3* (i, ii) or *cen5* (iii, iv). **(i)** WT (N = 3, n = 207) versus *pre9Δ* (N = 3, n = 423); **(ii)** WT (N = 2, n = 205) versus *rpt6-HA_3_* (N = 2, n = 127) **(iii)** mock (DMSO) (n = 156) versus MG132-treated (n = 155). Strains also carry an inducible *pGAL1-ndt80/” pdr5Δ/”*. **(iv)** Wild-type *PRE9* (N = 2, n = 237) versus *pre9Δ* (N = 2, n = 210) in a *spo11-yf* background. For segregation patterns in a *NDT80 pdr5Δ*/” strain treated with MG132 see **Fig. S2E**. For tetrad viabilities of WT, *pre9Δ,* and *rpt6-*HA_3_ see **Fig. S2C**. Schematic depiction of chromosomes 3 and 5 with centromeres and TetR-GFP binding sites represented to scale. cM (centimorgan) values indicate genetic distances between the locus carrying the *tetO* array and *cen3* or *cen5*, respectively. Data are presented as mean ± SD. Significance was determined by unpaired t-test with Welch’s correction and is represented by asterisks (*, p< 0.05; **, p< 0.01; ***, p< 0.001; ****, p< 0.0001; ns, not significant).

Examination of cells with one GFP-tagged homologue demonstrated normal sister chromatid cohesion during *pre9Δ* meiosis I, as indicated by GFP dots in only one nucleus of most binucleate cells (**Fig. 2B-i**). By contrast, during meiosis II, sister chromatids frequently failed to separate: tetrads containing GFP signals in only one instead of two nuclei were increased from 10 (± 3.4)% in wild type to 56 (± 8.3)% in *pre9Δ* (**Fig. 2A-i**; **Fig. 2B-ii**). Conversely, homologues disjoined normally during meiosis I in *pre9Δ*, as indicated by wild-type levels of homologue nondisjunction in binucleate cells carrying GFP on both copies of chromosome 5 (**Fig. 2B-iii**). We conclude that meiosis II sister chromatid nondisjunction (SNDJ) alone is responsible for the decreased viability of gametes, as neither meiosis I precocious separation of sister chromatids (PSSC) nor meiosis I homologue nondisjunction (HNDJ) are increased in absence of α3^Pre9^.

To better characterize the type of missegregation that occurs in *pre9Δ*, we considered three possibilities: (i) unequal segregation in which most chromosomes are pulled toward the same spindle pole; (ii) persistent linkages between sister chromatids always affecting both chromosomes of a homologue pair; and (iii) persistent linkages between sister chromatids affecting homologue pairs independently. These studies revealed that individual homologues, rather than both homologues or the entire genome complement, undergo sister chromatid non-disjunction during *pre9Δ* meiosis II. Accordingly, the four chromatin masses in tetranucleate cells exhibited comparable sizes and chromatin content, regardless of whether the GFP-tagged chromatid had segregated correctly or incorrectly (**Fig. 2A-i, right; Fig. 2C**). Thus, *pre9Δ* does not result in a general segregation bias to a subset of nuclei. Next, we analyzed whether all 4 chromosomes of a given homologue pair missegregated or whether the missegregation was limited to individual homologues. If both sister chromatids of a given homologue pair had failed to disjoin, tetrads with only two GFP-positive spores would predominate. Instead, we found that tetrads from cells carrying GFP tags on both homologues most often display GFP in three of four spores (∼two-thirds of cases), and less frequently in two of four spores (∼one-third; **Fig. 2B-iv**). Thus, meiosis II sister chromatid nondisjunction in *pre9Δ* occurs independently for each homologue, affecting ∼40% of cells in the population (**Fig. 2B-iv**).

Whereas α3^Pre9^ is critical for accurate meiosis II sister chromatid segregation, it is dispensable for chromosome segregation in meiosis I (above) and of sister chromatids during mitosis. Accordingly, in diploid cultures heterozygous for *cen5-*GFP, mitotic chromosome missegregation would generate subpopulations of trisomic and monosomic cells, the latter lacking a GFP signal if the marked chromosome is lost. Yet, no such signal loss was evident even after ∼30 consecutive rounds of mitotic cell divisions: among diploid cells arrested at G1, a GFP signal was detectable in > 90% of both *pre9Δ* and WT cells (n ≥ 125; **Fig. 2A-ii**). Furthermore, neither haploid nor diploid *pre9Δ* strains display mitotic growth defects, as evidenced by spore colony sizes indistinguishable from wild type, the absence of sectored *pre9Δ* spore segregants (**Fig. S2B-i**), and comparable colony growth of WT and *pre9Δ* diploids across a wide temperature range ^29^.

We conclude that α3^Pre9^, either alone or as part of the CP, is required specifically for sister chromatid disjunction in meiosis II, while arm cohesion during meiosis I and mitotic sister chromatid cohesion are removed normally in its absence. Thus, deletion of *α3* distinguishes proteasome functions in meiosis II chromosome segregation from those in meiosis I and mitosis. Furthermore, our data suggest that sister chromatid nondisjunction in *pre9Δ* may arise from stochastic persistence of linkages between sister chromatids in meiosis II, rather than from gross chromatin missegregation.

### α3^Pre9^ supports meiosis II sister chromatid disjunction through canonical 26S proteasome function

To determine whether the exquisite sensitivity of meiosis II chromosome segregation to α3^Pre9^ disruption reflects a canonical proteasome function or a subunit-specific activity, we examined meiosis II sister chromatid segregation in another proteasome subunit mutant. We extended with an HA_3_ tag the C-terminus of ATPase Rpt6 which is located at the base of the 19*S* regulatory particle. The C-terminus of Rpt6 protrudes into the intersubunit pocket between α3^Pre9^ and α2^Pre8^ of the CP α ring, which serves as a critical docking site for RP assembly ^7,32^. Like *pre9Δ*, the *rpt6-HA_3_*mutant exhibits wild-type vegetative growth at 30°C (**Fig. S2B-ii**), yet only 50% of cells complete the meiotic divisions, compared to ∼90% in wild type. Furthermore, meiotic divisions in *rpt6-HA_3_* in the *SPO11* background are delayed by ∼ 3 h, a qualitatively similar defect as observed in *pre9Δ* (23°C; **Fig. S2A**). *rpt6-HA_3_* cells heterozygous for *cen3*-GFP formed tetrads containing a fluorescent dot in one instead of two nuclei at high frequencies [∼39 (± 0.35)%], closely matching the *pre9Δ* phenotype [41 (± 3.2) % compared to ∼5% in wild type (**Fig. 2D-i**]. Consistent with increased chromosome missegregation, spore viability in the same meiotic cultures was also reduced, to ∼ 38 (± 3.5) % in *rpt6-HA_3_*and 48 (± 4.9) % in *pre9Δ*, compared to 95 (± 2.5) % in wild type (**Fig. S2C**). Tetrad viability patterns in proteasome mutant strains were consistent with aberrant sister chromatid segregation as indicated by a prominence of tetrads comprising 3, 2, or 1 viable spores, and not with homologue non-disjunction, where 2 and 0 viable spores would predominate (**Fig. S2C; Table S1**). Thus, like loss of CP component α3^Pre9^, alteration of its RP interaction partner Rpt6 selectively impairs meiosis II sister chromatid separation while leaving intact other proteasome functions, including those in mitotic and meiosis I chromosome segregation.

We also inhibited the proteasome chemically using MG132, a peptide aldehyde that blocks the chymotrypsin-like activity of the β5 proteasome subunit ^33^. To ensure intracellular drug retention, experiments were performed in a *pdr5Δ* background ^34^. Moreover, to maximize the proportion of cells in meiosis II during proteasome inhibition, MG132 was added to cultures three hours after release from prophase I arrest (**Fig. S2D-i**) ^35^. Cells that had passed the metaphase I-anaphase I transition before treatment either entered into or completed meiosis II, whereas cells still mononucleate at the time of treatment mostly arrested (**Fig. S2D-i,** dashed blue line). Importantly, ∼57% of the cells that completed meiosis II in the presence of MG132 exhibited sister chromatid non-disjunction, compared to only ∼8% in the mock-treated culture (**Fig. 2D-iii; Fig. S2D-i**; for details see Materials and Methods). Comparable results were obtained when meiotic cultures expressing *NDT80* under its own promoter were treated with MG132 at t = 7 h (**Fig. S2D-ii; Fig. S2E**).

Together, these findings show that canonical proteasome activities - both substrate engagement by the RP and proteolysis by the CP - are particularly critical for meiosis II sister chromatid segregation. While disrupting proteasome function through *pre9* (CP) or *rpt6* (RP) mutations does not block meiosis II chromosome segregation, accuracy of segregation critically depends on the proteasome function ensured by these subunits.

### Meiosis II defects in *pre9Δ* are unrelated to DSB-induced prophase I events

Because the proteasome is central to homologue pairing, recombination, and synapsis, as indicated by analysis under conditions that trigger meiotic prophase I arrest ^29^, we examined whether the meiosis II sister chromatid missegregation in proteasome mutants might be a secondary consequence of prophase I abnormalities. To test this, *pre9Δ* was combined with *spo11-yf* (above). Without Spo11-induced DSBs, homologues attach randomly to spindle poles, resulting in frequent meiosis I nondisjunction in a wild-type *PRE9* background, yet meiosis II sister chromatid segregation occurs normally (**Fig. 2D-iv**) ^36^. By contrast, *pre9Δ spo11-yf* still exhibited meiosis II sister chromatid non-disjunction in 44 (± 7.1) % of cells, very similar levels as those observed in the *pre9Δ SPO11* background (**Fig. 2D-iv**). Thus, defective meiosis II sister chromatid segregation in proteasome mutants is not a byproduct of a meiosis I proteasome role in DSB processing. Instead, independent of any roles in DSB-initiated events during meiosis I, a fully functional 26*S* proteasome is required specifically for meiosis II sister chromatid segregation.

### Proteomic analysis reveals an increase of cohesin protector Sgo1 in *pre9Δ*

To obtain unbiased insights into effects of the *pre9Δ* proteasome mutation on the abundance of proteins with functions in chromosome segregation, Tandem Mass Tag (TMT6plex) spectrometry was performed on synchronized meiotic cultures ^37^, comparing *pre9Δ* and wild-type *PRE9*. To minimize differences in the timing of meiosis II entry, experiments were carried out in a *spo11-yf* background. Synchronous meiosis was induced at 30°C and proteins were extracted from culture aliquots obtained at t = 6.5 h, a time when ∼50% of cells are at the binucleate stage in both wild-type *PRE9* and *pre9Δ* (**Fig. S3A**).

Among a total of 3,770 detected proteins, 597 proteins (∼16%) were differentially expressed in *pre9Δ* compared to wild-type *PRE9*, as indicated by inverse expression levels in samples from the two genotypes but strong correlation of expression levels within each genotype (**Fig. 3A, Fig. S3B; Table S2**). In a volcano plot of *PRE9*/*pre9Δ* values, about half of the 597 proteins are significantly (green) enriched in *pre9Δ* (left) and half are significantly enriched in wild-type *PRE9*, including Pre9 itself, as expected (**Fig. 3B**, right). Likely relating to a delay in *pre9Δ* in completing meiosis II, many proteins with roles in maturation of tetrads into asci are overrepresented in *PRE9* (**Fig. 3C**). Importantly, phosphatase PP2A recruiter shugoshin Sgo1, a centromeric factor with roles in controlling meiosis II sister chromatid disjunction, is overrepresented in *pre9Δ* (**Fig. 3B**). Given its critical role in ensuring cohesin protection until meiosis II ^17,22,24^, we further investigated the relationship between Sgo1^shugoshin^ and the Pre9-proteasome.

**Fig. 3.**
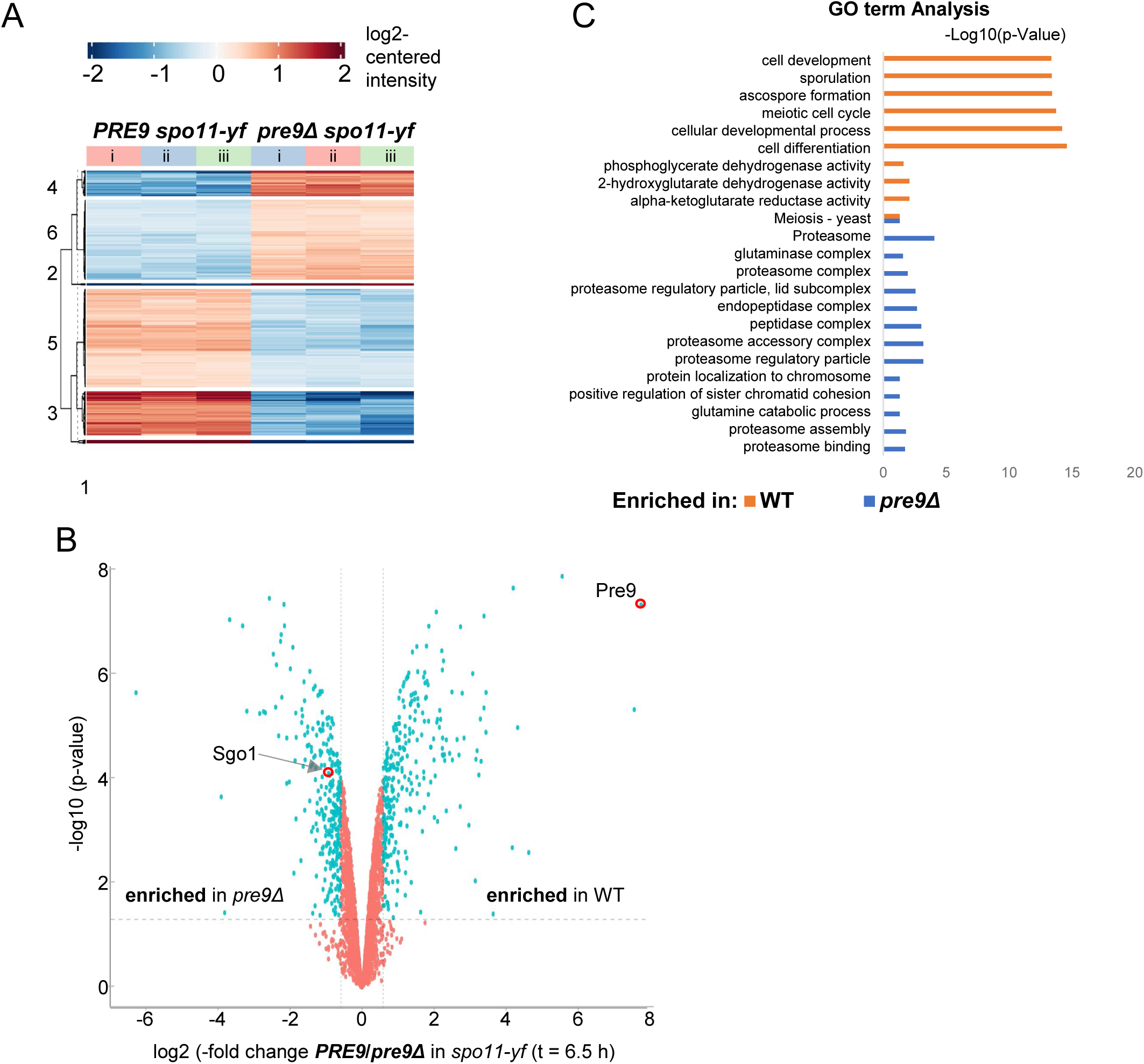
Mass spectrometry reveals increased abundance of meiosis I cohesin-protector Sgo1 in *pre9Δ*. (A) Cluster analysis indicates reproducibility of protein enrichment in triplicate samples (i, ii, & iii) of *PRE9* and *pre9Δ* protein samples in the *spo11-yf* background at t = 6.5 h at 30℃. For meiotic progression, see **Fig. S3A**. (B) Volcano plot of protein abundance in *PRE9* compared to *pre9Δ* in a *spo11-yf* background. Proteins over- or underrepresented in *pre9Δ* are on the left and right side of the plot, respectively. Cut offs are 1.5-fold. For a complete list of enriched and depleted proteins, see **Table S2**. (C) Gene ontology (GO) analysis of genes enriched or depleted in *pre9Δ* compared to *PRE9* in a *spo11-yf* background at t = 6.5 h.

### An α3^Pre9^-containing proteasome mediates removal of cohesin-protector Sgo1 from meiosis II kinetochores

Our proteomic analysis suggested that Sgo1 protein levels are increased in *pre9Δ* at least at one timepoint. Sgo1 recruits PP2A phosphatase to kinetochores during meiosis I, thereby countering Rec8 phosphorylation as prerequisite for separase cleavage (Introduction) ^22,24,38^. To examine *pre9Δ* effects on Sgo1 in more detail, we monitored its protein levels throughout synchronized meiosis. Western blot analysis of V5_3_-tagged Sgo1 (hereafter Sgo1-V5) in wild-type *PRE9* in a *spo11-yf* background revealed strong meiotic induction, with Sgo1 appearing at t = 6 h, followed by a broad peak and a return to minimum levels by t = 11 h, consistent with earlier findings (**Fig. 4A, B**) ^26^. In *pre9Δ*, Sgo1-V5 appeared with similar timing but reached higher peak levels before returning to wild-type levels by 11 h. Thus, cellular Sgo1 is elevated in *pre9*Δ throughout meiosis, corroborating the mass spectrometry analysis.

**Fig. 4.**
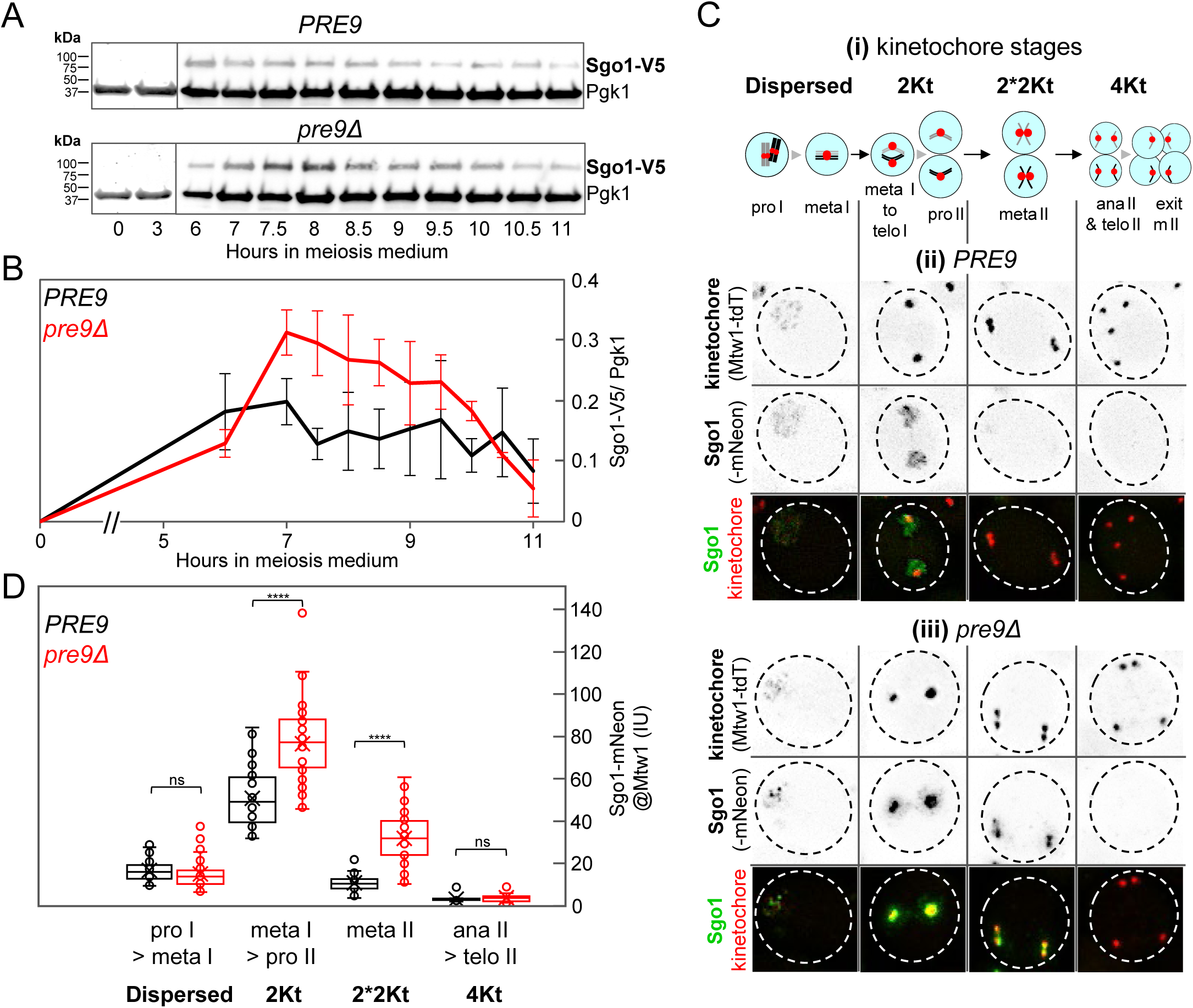
An α3^Pre9^-containing core proteasome mediates removal of cohesin protector Sgo1 from pericentromeric region. (A) Western blot analysis of cellular shugoshin levels (Sgo1-V5) in wild-type *PRE9* and *pre9Δ* meiotic cultures also carrying *spo11-yf*. Antibodies against V5 and PGK1 loading control were applied to membranes at the same time and blots from paired *PRE9* and *pre9Δ* cultures were analyzed in the same LI-COR scan. (B) Quantitative analysis of average cellular Sgo1-V5 levels in wild-type *PRE9* and *pre9Δ* cultures (n = 3). Levels are expressed as signal of Sgo1-V5 divided by Pgk1. Quantitation was carried out from the same LI-COR scan of *PRE9* and *pre9Δ.* Error bars indicate standard deviations. (C) Sgo1 has disappeared from kinetochores by metaphase II in wild-type *PRE9*, but remains associated with metaphase II kinetochores at substantial levels in *pre9Δ*. **(i)** Schematic depiction of meiotic cell division stages. Kinetochores are in red, one representative homologue pair in black and grey, nuclei in light blue. Kinetochore (Mtw1-tdTomato) and shugoshin (Sgo1-mNeonGreen) fluorescence at the indicated stages in representative fixated **(ii)** wild-type *PRE9* and **(iii)** *pre9Δ* cells. Dashed lines indicate cell wall. Size bars are 1 μm. (D) Quantitative analysis of Sgo1 fluorescence intensity in *PRE9* and *pre9Δ* cells from prophase I until completion of meiosis II. See Fig. 4C**-i** for schematic representation of the four stages. Number of nuclei is ≥ 15 for each stage. Box-and-whisker plots show the median (center line), the interquartile range (box), and whiskers extending to 1.5 × interquartile range, with individual outliers shown as points. Statistical significance is represented by asterisks (*, p < 0.05; **, p < 0.01; ***, p < 0.001; ****, p < 0.0001; ns, not significant).

To determine whether elevated Sgo1 resides at kinetochores, we quantitated in intact cells the fluorescence intensity of Sgo1-mNeonGreen (Sgo1-mNeon hereafter) signal co-localizing with the kinetochore marker Mtw1-tdT at the same four cytological stages analyzed for meiotic progression (see **Fig. 4C-i**). In wild type (*PRE9 SPO11*), Sgo1 localizes to dispersed prophase I kinetochores at low levels (**Fig. 4C-ii**), consistent with earlier findings ^26^. At the 2Kt stage, strong Sgo1 signal surrounds and overlaps with kinetochores, likely reflecting pericentromeric enrichment. In metaphase II (2*2Kt), Sgo1 signal is faint, even though some remnants are detectable on chromatin spreads ^17,18^, becoming undetectable in all cells by the 4Kt stage (**Fig. 4C-ii**). In *pre9Δ SPO11* during prophase I, Sgo1 intensity and patterns are similar to those in wild-type *PRE9*. In contrast, at the 2Kt stage, Sgo1 staining is more intense and more concentrated at kinetochores of *pre9Δ* cells than in wild type, and this intense staining persists through metaphase II (2*2Kt), when Sgo1 levels associated with kinetochore pairs are at least 3-fold higher in *pre9Δ* than in wild type (**Fig. 4C-iii; Fig. 4D**). The Sgo1 signal typically is closely associated or partially overlaps with the kinetochore signal in metaphase II cells, potentially indicating association with the pericentromeric region (**Fig. 4C-iii; Fig. 4D**; n > 35 in each class). As in wild type, by the 4Kt stage Sgo1 is absent from kinetochores in *pre9Δ*.

We conclude that an α3^Pre9^-containing proteasome limits the levels of Sgo1 at kinetochores and is required for its timely removal from kinetochores. Importantly, Sgo1 remains prominent at *pre9Δ* metaphase II kinetochores, a stage when it is undetectable in intact wild-type cells. Its increased intensity further indicates that persistence of Sgo1 is not a trivial consequence of delayed meiotic progression which would increase the frequency of a given stage, but not the levels of protein at kinetochores.

### An α3^Pre9^-proteasome mediates timely Rec8 removal from pericentromeric regions

If the prolonged kinetochore association of Sgo1 in *pre9Δ* anchors PP2A, pericentromeric Rec8 would be expected to persist as well, since Sgo1-PP2A dephosphorylation shields Rec8 from separase cleavage. To examine this possibility, cellular Rec8 protein levels were monitored using V5-tagged Rec8 in a *spo11-yf* background. In wild-type *PRE9*, full length Rec8 becomes detectable at t = 3 h, reaches peak levels around t = 6 h, and has returned to very low levels by t = 10 h (**Fig. 5A, B**). In *pre9Δ*, Rec8 initially appears with levels and kinetics similar to those observed in wild type, yet it persists at higher levels and for a longer time compared to *PRE9*, even though no significant increase was detected by mass spectrometry (**Fig. 5A,B**; **Table S2**).

**Fig. 5.**
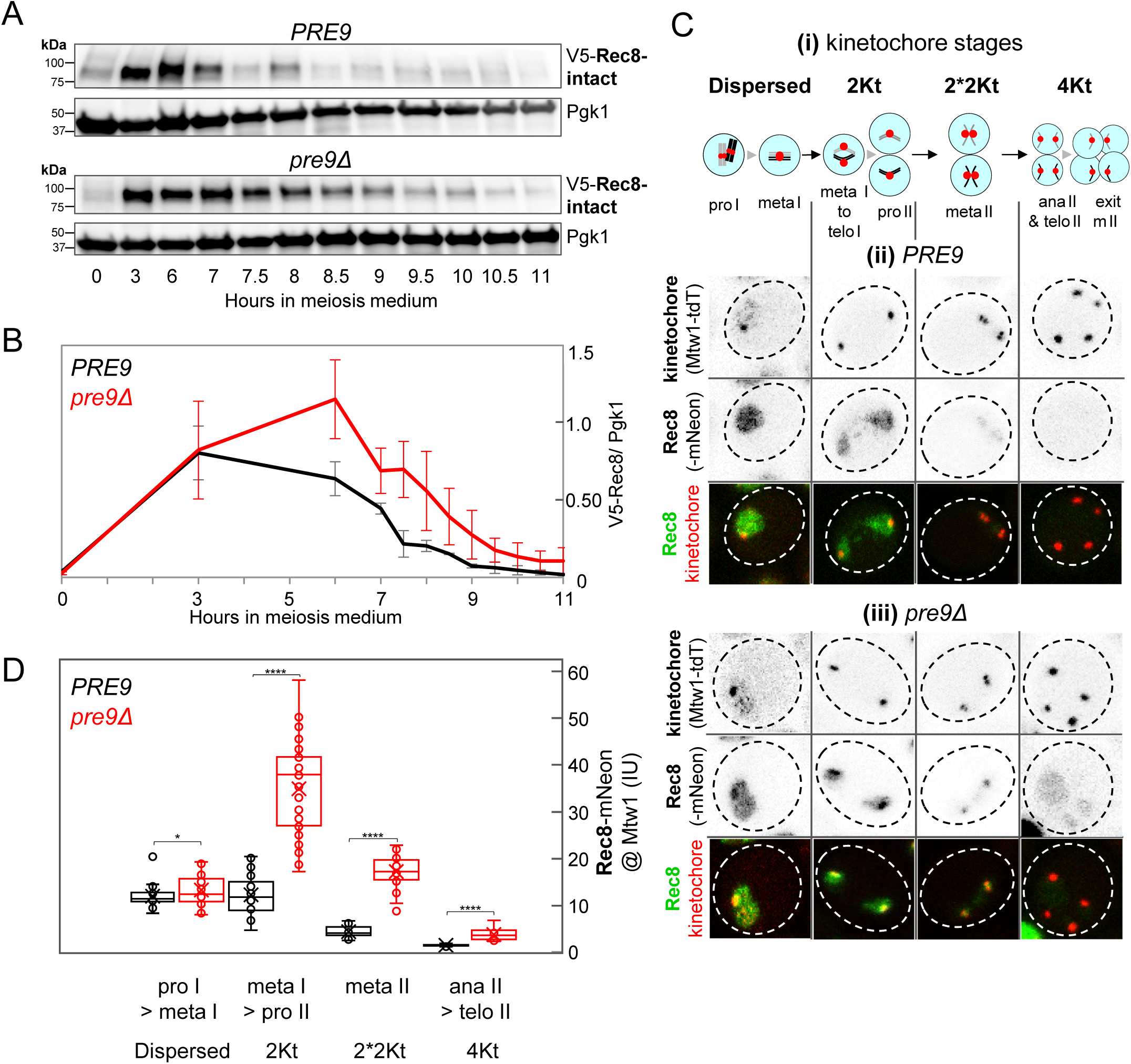
An α3^Pre9^-containing core proteasome mediates removal of cohesin Rec8 from kinetochores. (A) Western blot analysis of cellular Rec8 levels (V5-Rec8) in synchronized wild-type *PRE9* and *pre9Δ* cultures also carrying *spo11-yf*. An antibody against phosphoglycerate kinase 1 (Pgk1) was used as loading control. (B) Quantitative analysis of full length V5-Rec8 levels in *PRE9* and *pre9Δ* cultures. Levels are expressed as signal of V5-Rec8 divided by Pgk1. (C) Most Rec8 has disappeared by metaphase II in wild-type *PRE9* but in *pre9Δ* remains associated with kinetochores into metaphase II at substantial levels. **(i)** Schematic depiction of meiotic cell division stages between prophase I and completion of meiosis II. **(ii)** Kinetochore (Mtw1-tdTomato) and Rec8 (Rec8-mNeon) fluorescence at the indicated stages in representative fixated *PRE9* and **(iii)** *pre9Δ* cells. Dashed lines indicate cell wall. Size bars are 1 μm. (D) Quantitative analysis of fluorescence intensity of Rec8 at Mtw1-marked kinetochores in wild-type *PRE9* and *pre9Δ* cells from prophase I to the end of meiosis II. See Fig. 5C**-i** for schematic representation of the four stages. Number of nuclei is ≥15 for each stage. Significance was determined by unpaired t-test with Welch’s correction. Statistical significance is represented by asterisks (*, p < 0.05; **, p < 0.01; ***, p < 0.001; ****, p < 0.0001; ns, not significant).

To ask whether *pre9Δ* affects the levels of Rec8 at centromeres, we measured the fluorescence intensity of mNeonGreen-tagged Rec8 (hereafter Rec8-mNeon) co-localizing with Mtw1-tdT from prophase I through completion of meiosis II, staging intact cells based on kinetochore status (above; **Fig. 5C-i**). During wild-type *PRE9* meiosis, Rec8 is prominently detected throughout the nucleus in prophase I (dispersed stage), converging to and around kinetochores until prophase II (2Kt stage; **Fig. 5C-ii**). By metaphase II (2*2Kt stage), the Rec8 signal is reduced approximately 3-fold compared to the 2Kt stage (**Fig. 5C-ii; 5D**). Consistent with earlier reports, Rec8 is undetectable after metaphase II (4Kt stage; **Fig. 5C-ii**) ^16,26^.

In *pre9Δ* prophase I, Rec8 localizes throughout the nucleus with some enrichment at kinetochores similarly as in wild type, although Rec8 levels are marginally increased (**Fig. 5C-iii; 5D**). In contrast, in 2Kt cells, Rec8 intensity is ∼4-fold higher in *pre9Δ* than in wild-type, with Rec8 forming strong foci at the two kinetochores in addition to halos surrounding them. In metaphase II cells in *pre9Δ*, the Rec8 signal remains 4-fold increased, while little Rec8 remains associated with kinetochores in wild-type *PRE9* (**Fig. 5C-iii**). Remarkably, a single Rec8 focus is now located between each of the two kinetochore signal pairs (2*2Kt), unlike Sgo1 which exhibits a more varied association with kinetochore pairs (compare third frames in **Fig. 4C-iii** and **5C-iii**). Finally, even in post-metaphase II cells (4Kt), the Rec8 signal, though substantially decreased, remains significantly stronger in *pre9Δ* compared to wild-type *PRE9* (**Fig. 5C, D**).

We conclude that in meiosis II, both Sgo1 and the kleisin Rec8 are significantly enriched at kinetochores in *pre9Δ* cells relative to wild type, and that both proteins persist into later stages of the cell cycle. The presence of Rec8 between metaphase II kinetochore pairs in *pre9Δ* cells is consistent with kinetochores under tension that fail to separate due to persistent Rec8-mediated cohesion. Moreover, the continued presence of low levels of Rec8 in the centromeric region in *pre9Δ* beyond metaphase II supports the idea that persisting cohesion is responsible for failed separation of a subset of sister chromatids in meiosis II.

### The α3^Pre9^-proteasome coordinately controls Rec8 displacement and metaphase II completion

To examine in more detail Rec8 dynamics during meiosis II sister chromatid separation, we performed live-cell imaging of Rec8-mNeon and kinetochore marker Mtw1-tdT, again in a *spo11-yf* background to ensure similar timing of meiosis I divisions. Images were acquired in 10 minute intervals starting prior to metaphase II entry, which was identified by the transition from the 2Kt to the 2*2Kt stage and defined as t = 0 h in all genotypes, and ending after completion of anaphase II, which was identified by separation of both kinetochore signals (> 1 μm apart; 4Kt; see schematics in **Fig. 6A-i, ii**).

**Fig. 6.**
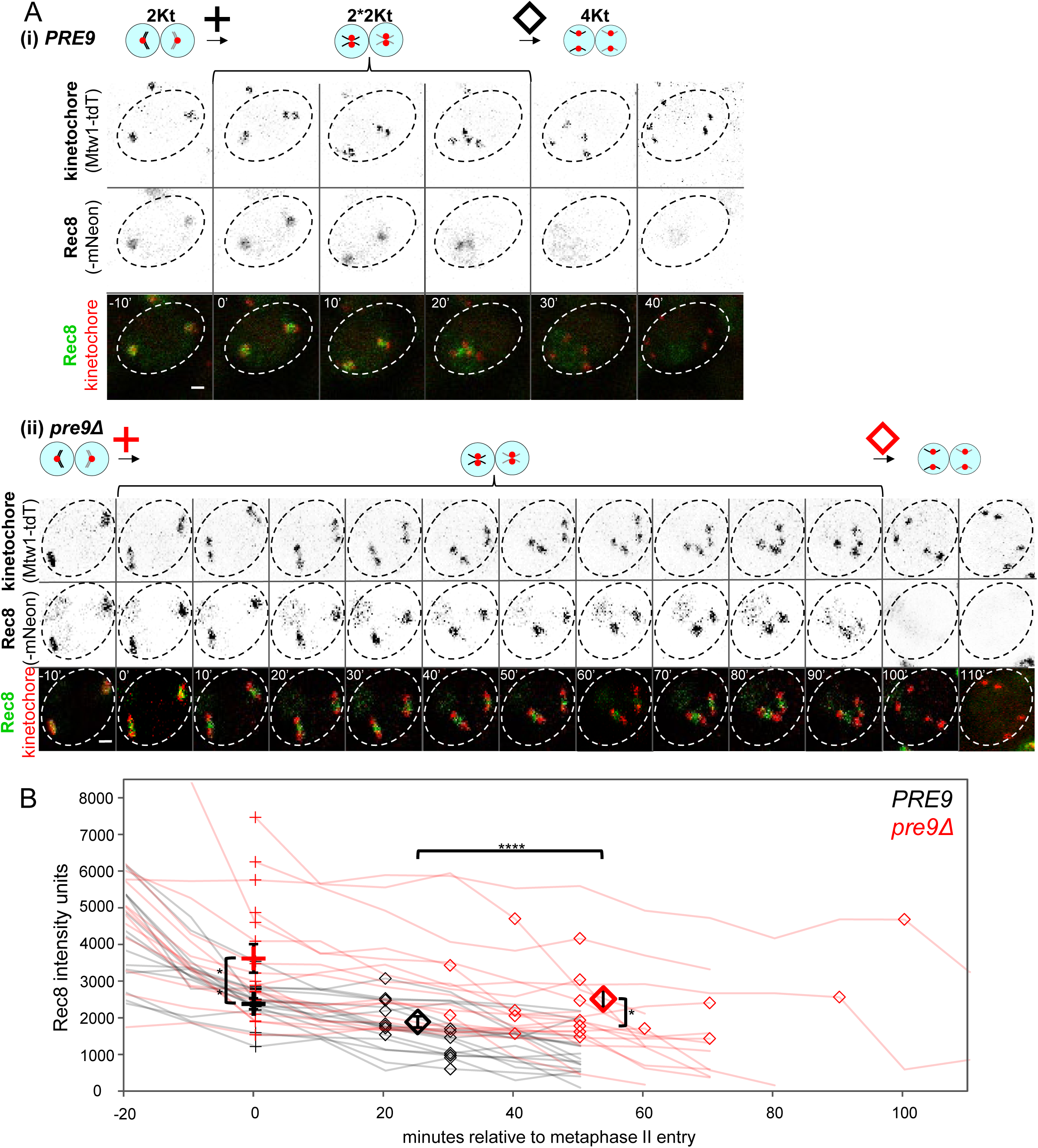
Rec8 is retained at higher levels and persists in *pre9Δ* relative to *PRE9* during the metaphase II–anaphase II transition. (A) Time lapse imaging of Rec8-mNeon association with Mtw1-marked kinetochores in representative (i) wild-type *PRE9* and (ii) *pre9Δ* cells from metaphase II entry to exit. Images were collected in 10-minute intervals. Brackets indicate the time from entry into metaphase II, marked by a cross [t = 0 min; indicated by the first frame with 2 kinetochore pairs per cell] to entry into anaphase II, marked by a diamond [4 kinetochore signals separated by more than 1 µm]. The final images show cells at or beyond the anaphase II stage. Size bars are 1 μm. (B) Quantitative analyses of Rec8-mNeon signal intensity at Mtw1-marked kinetochores in wild-type *PRE9* (black lines and symbols) and *pre9Δ* (red lines and symbols). Rec8 fluorescence intensities were recorded from live cell imaging frames taken at 10 min intervals. Small crosses indicate the time and Rec8 levels when a given cell has entered metaphase II (the 2*2Kt stage), set to t = 0 for all cells (except for three *pre9Δ* nuclei, for which pre-metaphase II measurements are missing). Small diamonds indicate the time and Rec8 intensity of the same cells in the last frame prior to anaphase II. Large crosses indicate average Rec8 levels in the first metaphase II frame, large diamonds indicate average Rec8 levels at the time of exit from metaphase II. Error bars are SD of Rec8 intensity levels at the respective stages. Significance is indicated by asterisks (*, p< 0.05; **, p< 0.01; ***, p< 0.001; ****, p< 0.0001; ns, not significant).

In wild-type *PRE9*, Rec8-mNeon was detectable as a cloud of puncta bridging paired kinetochores until the last metaphase II frame and declined to background levels upon anaphase II entry, consistent with earlier findings (**Fig. 6A-i**) ^16,26^. Two major differences were apparent in *pre9Δ* compared to *PRE9*: First, Rec8 levels in *pre9Δ* cells were increased 1.6-fold at metaphase II onset (**Fig. 6B**) and 1.4-fold at anaphase II onset. Second, paired and closely juxtaposed kinetochore signals (2*2Kt) persisted in *pre9Δ* cells for 55 (± 22) min (n = 18) compared to 25 ± 5 min (n = 17) in *PRE9*, indicating a ∼two-fold increase in metaphase II duration (**Fig. 6B; Fig. S4**). Additionally, paired metaphase II kinetochores (2*2Kt) in *pre9Δ* tended to be spaced farther apart over multiple consecutive frames, with bridging Rec8 appearing elongated instead of punctate as in wild-type *PRE9* (**Fig. 6A-ii**). The increased Rec8 intensity was more pronounced in fixed cells compared to live cells, likely due to selection bias, resulting from the fact that our live-cell analysis only included cells that progressed through the 2Kt→2*2Kt→4Kt transition within the imaging time frame, while cells that failed to progress to the 4Kt stage were excluded from analysis. Consistent with this interpretation, prolonged metaphase II duration among live *pre9Δ* cells correlated with elevated Rec8 intensity (**Fig. 6B**).

Thus, in the absence of the proteasome α3^Pre9^ subunit, cohesin Rec8 levels are higher at kinetochores, and the duration of metaphase II is at least doubled. Notably, both Rec8 intensity and metaphase II duration were more variable in *pre9Δ* than in wild-type *PRE9*, indicating heterogeneity of these effects in *pre9Δ*. This is consistent with observations from fixed-cell analyses (**Fig. 5C, D**) and may explain why only a subset of chromosomes fail to separate during meiosis II in *pre9Δ*.

### Induced Rec8 destabilization partially bypasses the α3^Pre9^-proteasome requirement during meiosis II

To test whether α3^Pre9^ affects meiosis II sister chromatid segregation through a role in Rec8 removal and/or cleavage, we monitored the appearance of separase-cleaved V5-Rec8 ^13,39^, again in a *spo11-yf* background to ensure comparable timing of meiosis II entry (**Fig. 7A**). In wild-type *PRE9*, cleavage of full length V5-Rec8 by separase generates two similarly sized N-terminal fragments which became detectable as a single band at ∼t = 8 h, reached levels equal to those of full-length Rec8 by ∼t = 10 h, and became the dominant band thereafter (**Fig. 7A-i, ii;** see diagram in **Fig. 7C**). In *pre9Δ*, the Rec8 separase cleavage product appeared at roughly the same time as in wild type, but its levels remained below 20% of the total Rec8 signal even at late time points – long after meiosis I was completed and arm cohesin is normally cleaved (**Fig. 7A-iii**). Thus, in the *pre9Δ* mutant, full length Rec8 persists and its cleavage remains incomplete.

**Fig. 7.**
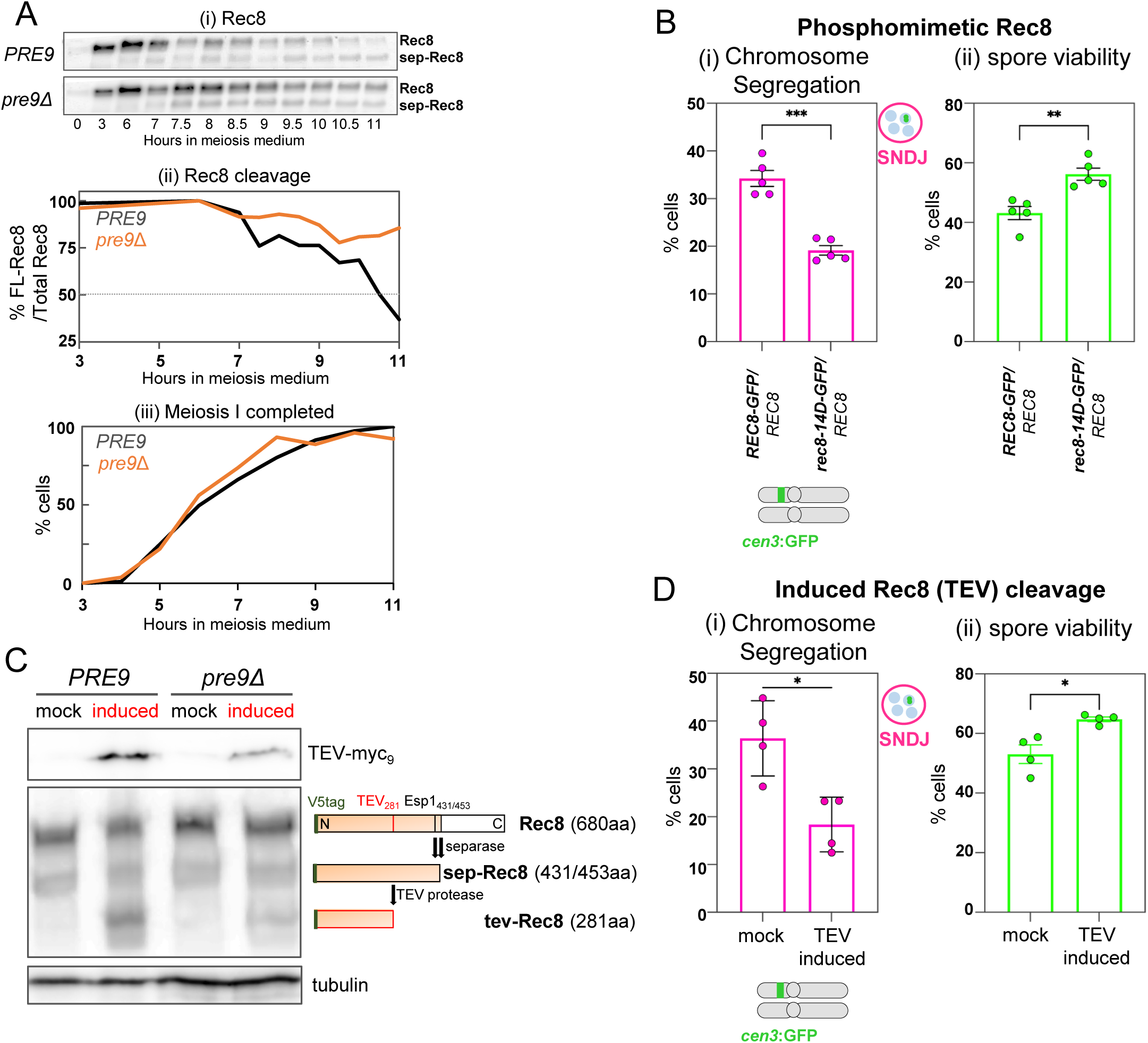
Phosphomimetic Rec8 and induced Rec8 cleavage partially rescue sister chromatid disjunction and spore viability in an α3^Pre9^-deficient proteasome mutant. (A) **(i)** Western blot analysis in wild-type *PRE9* and *pre9Δ* in a *spo11-yf* background of full-length V5-Rec8 (76 kD) and N-terminal V5-Rec8 separase cleavage product “sep-Rec8” (∼ 50 kD; see diagram in Fig. 7C). **(ii)** Quantitative analysis of cellular full-length Rec8 levels as a percentage of total (i.e. full length + cleaved) V5-Rec8. **(iii)** Meiotic nuclear divisions for the cultures analyzed for V5-Rec8 separase cleavage. (B) Heterozygosity for the phosphomimetic *rec8-14D-GFP/ REC8* allele in a *pre9Δ* mutant **(i)** partially restores normal meiosis II sister chromatid disjunction and **(ii)** improves spore viability. Segregation of heterozygous *cen3*-GFP and spore viability were determined following incubation in meiosis medium for 24 hours at 23°C. For meiotic divisions, see **Fig. S5A**. Data are presented as mean ± SEM. Significance here and in Fig. 7D was determined by unpaired t-test with Welch’s correction; p-values are indicated by asterisks (*, p< 0.05; **, p< 0.01; ***, p< 0.001; ****, p< 0.0001; ns, not significant). (C) Western blot analysis of Rec8 cleavage by TEV protease. **Top panel:** Detection of C-terminally myc_9_-tagged TEV protease expressed under control of the *pGAL1* promoter upon induction with β-estradiol (see text). **Middle:** N-terminally V5-tagged Rec8 carrying a TEV protease recognition site at aa281. Detection in *PRE9* and *pre9Δ* meiotic cultures of uncleaved (Rec8), separase-cleaved (sep-Rec8) and TEV-cleaved Rec8 (tev-Rec8). The expected lengths of Rec8 species excluding the 14 aa V5 tag are indicated. **Bottom:** α-tubulin loading control. (D) Induction in a *pre9Δ* mutant of ectopic Rec8 cleavage through induced TEV protease at the time of meiosis II **(i)** partially restores normal meiosis II sister disjunction and **(ii)** improves spore viability (n = 4). Data are presented as mean ± SEM. For meiotic divisions, see **Fig. S5C**. For significance see Fig. 7B.

One mechanism by which the proteasome may contribute to meiosis II Rec8 cleavage is enhanced dephosphorylation of pericentromeric Rec8, potentially due to continued association of the phosphatase tether Sgo1 with centromeres and/or kinetochores (**Fig. 4**). Rec8 is phosphorylated at ≥ 26 positions, 14 of which are sufficient to induce precocious meiosis I sister chromatid segregation ^14,15^. We therefore introduced one allele of phosphomimetic *rec8-14D* while retaining one wild-type allele to avoid the precocious meiosis I separation of sister chromatids in homozygous *rec8-14D-GFP*, reported previously and confirmed in our hands ^15^ [**Table S1**]. Remarkably, heterozygous *rec8-14D-GFP/ REC8* significantly reduced sister chromatid nondisjunction in the *pre9Δ* background from 33 (± 3.3) % to 19 (± 2.0) % (**Fig. 7B-i**). Spore viability was also increased from 43 (± 2.8) % to 56 (± 3.2) % (**Fig. 7B-ii; Fig. S5B-ii**). Conversely, in the wild-type *PRE9* background, *rec8-14D-GFP*/ *REC8* exhibited spore viability indistinguishable from *rec8-GFP*/ *REC8* controls, further excluding the possibility that *rec8-14D-GFP*/ *REC8* results in increased PSSC during meiosis I (≥ 90%; **Table S1**).

If inefficient Rec8 phosphorylation in *pre9Δ* contributes to continued sister chromatid cohesion into anaphase II, cleaving Rec8 independently of phosphorylation status and separase activity should rescue sister chromatid separation even with defective proteasome function. We tested this idea using V5-Rec8 harboring an engineered TEV protease recognition site at aa281, combined with a system allowing inducible expression of TEV protease during meiosis (**Fig. 7C**) ^35,39^.

In a control experiment, induction at t = 5 h resulted in strong TEV protease expression in both wild-type *PRE9* and *pre9Δ* (**Fig. 7C-top**). A ∼35 kDa V5-Rec8 fragment, expected following TEV cleavage at aa281 (tev-Rec8), was detectable only in induced, but not in mock cultures (**Fig. 7C-middle**). By contrast, the ∼55 kDa V5-Rec8 fragment expected following cleavage by separase (sep-Rec8) was detected in both induced and mock cultures. Next, TEV expression was induced in *pre9Δ SPO11* at t = 9.5 h, when ≥ 40% of cells had completed meiosis I, thereby minimizing induced Rec8 cleavage during meiosis I (**Fig. S5C**). Remarkably, TEV-mediated Rec8 cleavage in *pre9Δ* resulted in a two-fold reduction in chromosome 5 sister non-disjunction, from 36 (± 6.8) % in mock cultures to 18 (± 4.9) % in induced cultures (**Fig. 7D-i**). Spore viability in the same cultures increased to 67 (± 1.1)% in induced cultures from 53 (± 5.4)% in mock cultures (p = 0.0287; n = 4; **Fig. 7D-ii**).

Albeit partial, the rescue of meiosis II sister chromatid segregation and spore viability upon TEV protease expression in *pre9Δ* is remarkable for several reasons. First, TEV-mediated rescue of chromosome segregation is likely limited to a subset of cells, since a substantial fraction may have already progressed past metaphase II before TEV expression, too late for rescue. Second, many *pre9Δ* cells retain elevated levels of Rec8 at kinetochores during metaphase II (**Fig. 5D**), which would be expected to impede cleavage and rescue of segregation. Third, TEV-dependent Rec8 cleavage likely is inefficient in *pre9Δ*. Notably, TEV induction in the wild-type cells leaves chromosome segregation essentially unaffected, as indicated by high spore viability (**Table S1**), suggesting that TEV cleaves Rec8 incompletely even under otherwise normal conditions. Finally, the partial rescue of *pre9Δ* chromosome segregation defects through TEV protease cleavage or phosphomimetic Rec8 could reflect technical limitations, including limited TEV protease activity and the presence of dephosphorylatable Rec8 in heterozygous *rec8-14D*/ *REC8* cells. Alternatively, it raises the possibility that the proteasome also promotes Rec8 deprotection through additional mechanisms (Discussion).

Together, these findings support a model in which the 26*S* proteasome, acting through a function dependent on its α3^Pre9^ subunit, promotes increased levels of pericentromeric Rec8 removal and thereby subsequent cleavage by separase. This activity involves removal of Sgo1 from kinetochores and possibly of additional antagonists of Rec8 phosphorylation. Notably, this requirement is specific to pericentromeric cohesion during meiosis II, as a global defect in Rec8 phosphorylation should result in meiosis I chromosome segregation defects ^14,15^, a defect not observed in *pre9Δ*. Importantly, the role of the proteasome in Rec8 cleavage identified here is likely distinct from its role in securin degradation, since *pre9Δ* cells cleave sufficient cohesin along chromosome arms to support normal homologue segregation during meiosis I and pericentromeric cohesion to allow substantial sister chromatid disjunction during meiosis II (Discussion).

### The proteasome core particle associates with kinetochores during meiosis II

To investigate how the proteasome might target and eliminate a centromere-associated protein such as Sgo1, we examined proteasome localization throughout meiosis using Mtw1-tdT as a kinetochore marker together with GFP-tagged core particle (α5^Pup2^) or regulatory particle (Rpn12) subunits ^29^.

In intact meiotic cells, from prophase I to telophase II, both CP (top) and RP (bottom) were substantially enriched in the nucleus compared to the cytoplasm (**Fig. 8A**), consistent with earlier findings in *S. pombe* ^40^. To determine whether the proteasome associates with chromatin after meiosis I, we squashed wild-type protoplasts from the t = 7 h time point in presence of detergent to remove much of the cytoplasm and nucleoplasm, then stained with antibodies against CP and kinetochores. Following this treatment, CP signal was largely absent from chromatin itself, although a halo of signals remained around the chromatin of binucleate meiosis II cells, likely derived from residual nucleo- and/or cytoplasm. A notable exception was a substantial proteasome aggregate consistently detected at the periphery of chromatin, as illustrated by three representative meiosis II cells (**Fig. 8B**). In 78% of binucleate cells, this CP signal overlapped with or was closely associated with kinetochore bundles in both nuclei, and with one of the two nuclei in an additional 19% (n = 27).

**Fig. 8.**
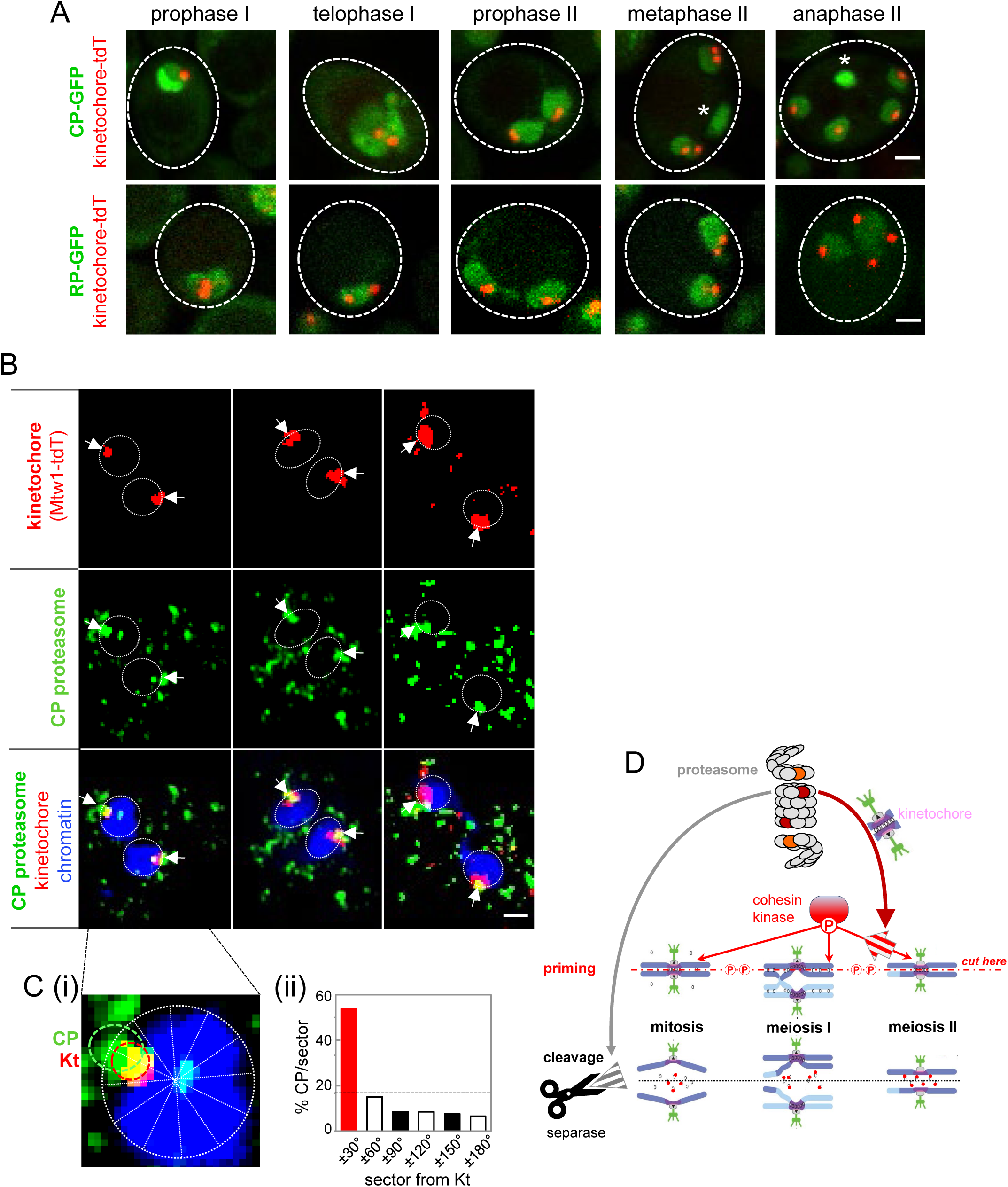
Localization of the proteasome during meiosis. (A) Core particle subunit á5^Pup2^-GFP (top) and regulatory particle subunit Rpn12-GFP (bottom) predominantly localize to the nucleoplasm in intact meiotic cells at the indicated meiosis I and meiosis II stages. Channels show GFP together with kinetochore (Mtw1-tdTomato) fluorescence used for staging (see **Fig. S1**). Asterisks: An additional GFP-fluorescing aggregate devoid of a kinetochore signal is frequently detected in meiosis II cells. Dashed lines indicate cell walls. Size bars are 1 μm. (B) Three representative binucleate, surface-squashed meiosis II cells immunodecorated with antibodies directed against GFP and tdTomato. α5^Pup2^-GFP and Mtw1-tdTomato are shown individually as well as together with DAPI-stained chromatin in the merge. Dashed lines mark nuclei. Size bar is 1 μm. (C) **(i)** Enlarged excerpt of a nucleus shown in (B), marking proteasome CP (green dotted line), the kinetochore (red line) and the 12 sectors with 0° defined by the centroid of the kinetochore signal. **(ii)** Histogram showing the frequency of proteasome α5^Pup2^-GFP signals in 30° bins centered on the kinetochore, with signals from both directions pooled into each bin. The two sectors flanking the kinetochore centroid are shown in red. The dashed line indicates the expected CP occupancy for random distribution across the 6 pooled sectors (16.6%). (D) Model of proteasome pathways controlling sister chromatid cohesion during the three types of cell division. Proteasome components α3^Pre9^ (CP) and Rpt6 (RP) are highlighted in red and orange, respectively. For details see Discussion.

To distinguish the *bona-fide* kinetochore-associated proteasome signal from incidental colocalization, we divided a circle encompassing each nucleus and 0.5 μm of its periphery into 12 equal 30° sectors, with the centroid of the kinetochore signal defined as 0° (**Fig. 8c-i**). On average, each nuclear area contained 1.96 CP foci. Of the total of 106 CP foci, 54% fell within the two sectors (-30° to +30°) containing the kinetochore, compared to an average of 9 (± 3) % [SD] in each of the remaining five equidistant sector pairs (**Fig. 8C-ii**), indicating a highly significant deviation from chance (p < 0.0001). We conclude that during wild-type meiosis, the proteasome core particle is spatially associated with kinetochores and/or the pericentromeric region at the stage when Sgo1 and Rec8 are displaced from kinetochores.

## Discussion

The proteasome plays a well-established role in chromosome segregation, driving securin degradation and consequent separase activation. Here, we have discovered that the proteasome also promotes cohesin cleavage along the second, parallel branch of this pathway. This second branch is specifically required for sister chromatid segregation in meiosis II (**Fig. 8D**). It is defined by its dependence on an α3^Pre9^-containing proteasome core particle which facilitates removal of the cohesin protector shugoshin Sgo1 and cohesin subunit Rec8 from centromeric regions. This function is associated with a spatially restricted pool of centromeric proteasomes. By regulating cohesin cleavage not only through separase activation but also by promoting Rec8 deprotection, the proteasome likely provides a fail-safe mechanism to counter residual separase activity carried over from meiosis I.

### Cohesin removal through a securin-independent proteasome pathway mediates accurate meiosis II sister chromatid disjunction

The current work reveals a stage-specific proteasome function in removing centromeric cohesion as a prerequisite for accurate meiosis II sister chromatid segregation, one that is separable from its established role in securin degradation. Several observations underscore that sustained cohesion in proteasome mutants prevents normal meiosis II sister chromatid disjunction. First, in cells lacking CP subunit α3^Pre9^, Rec8 persists at kinetochores at elevated levels as cells progress into anaphase II, well beyond its normal window of removal (**Fig. 5D**). Second, Rec8 cleavage is markedly delayed, extending into time points when cells have entered meiosis II (**Fig. 7A**). Third, induced Rec8 deprotection or cleavage partially restores sister chromatid disjunction in *pre9Δ* (**Fig. 7B-D**). Fourth, the prolonged presence of bi-oriented sister chromatids in *pre9Δ* metaphase II cells indicates that spindle forces act against unresolved cohesion (**Fig. 6B**). Together, these observations suggest that some cohesin rings remain intact in mutant cells resulting in meiosis II sister chromatid nondisjunction and formation of aneuploid gametes. Importantly, proteasome mutants lacking α3^Pre9^ or expressing separation-of-function allele *rpt6-HA_3_* are functionally intact for cohesin removal along chromosome arms, as chromosomes segregate normally during mitosis and meiosis I. Thus, these mutants remain intact for efficient securin degradation, but are specifically impaired for cohesin removal from pericentromeric regions in meiosis II.

Lack of cohesin cleavage has well-documented consequences for chromosome behavior resulting in metaphase arrest in mitosis, meiosis I, and meiosis II ^13,16,41,42^. Unlike these absolute impairments to chromosome segregation, meiosis II in *pre9Δ* and *rpt6-HA_3_* proteasome mutants eventually progresses, yet sister chromatids frequently fail to become separated from each other. Thus, cohesin persistence on only a few chromosomes may result in chromosome missegregation, but it is insufficient for entirely blocking meiotic divisions, indicating a lack of coordination between sister chromatid separation and meiosis II metaphase-to-anaphase progression.

### Proteasome-dependent shugoshin removal contributes to centromeric Rec8 deprotection

Stage-appropriate dynamics of shugoshin at kinetochores are critical for both mitotic ^43^ and meiotic chromosome segregation ^16–18,22,44,45^. Our findings implicate the ubiquitin-proteasome system in the timely removal of shugoshin from meiosis II kinetochores, with a fully intact proteasome required both to limit shugoshin levels and its prompt clearance. This defects is associated with impaired Rec8 cleavage, suggesting that the proteasome contributes directly to Sgo1 displacement during meiosis II.

Previously proposed mechanisms for relieving shugoshin/PP2A-mediated protection of kleisin cleavage include physical separation of shugoshin from kleisin ^25^, removal of a PP2A inhibitor ^46^, and APC/C-mediated polyubiquitination and ensuing proteasomal degradation of shugoshin ^16,26^. However, mutating predicted Sgo1 ubiquitination sites does not substantially increase sister chromatid non-disjunction, although it increases Sgo1 stability and delays its removal from kinetochores ^26^. In fact, reduced spore viability in cells expressing a non-ubiquitinatable Sgo1 allele requires its additional overexpression ^16,47^ suggesting that impaired ubiquitination alone is not sufficient to compromise segregation. Notably, whereas the proteasome mutant analyzed here causes pericentromeric Sgo1 accumulation (**Fig. 4**), the non-degradable Sgo1 alleles described previously persist at kinetochores without accumulating ^16,26^.

One explanation for these findings is that the proteasome mediates Rec8 deprotection at least in part through ubiquitination-independent displacement of Sgo1. This mechanism could involve recognition of an N-terminal degron, intrinsically disordered regions ^48^ and/or proteasome-driven chromatin remodeling ^46^ resulting in separation of Sgo1 from Rec8. Alternatively, and not mutually exclusively, the proteasome may facilitate meiosis II sister chromatid segregation by degrading additional antagonists of centromeric Rec8 removal.

### Proteasome-mediated control of cohesin cleavage via parallel pathways provides a fail-safe in meiosis II

Our observations reveal that the proteasome controls meiosis II sister chromatid segregation through removal of factors that control Rec8 cleavage, a function separable from securin degradation. Thus, the proteasome governs the two rounds of meiotic chromosome segregation through two parallel pathways: meiosis I is triggered by securin degradation and ensuing separase activation, while meiosis II additionally requires Rec8 deprotection via the proteasome-mediated removal of its phospho-regulators.

What is the physiological rationale for cohesin removal in meiosis II via two parallel pathways controlled by the proteasome? At the meiosis I metaphase-anaphase transition, robust separase activity is unleashed through APC-initiated securin degradation, enabling cohesin cleavage along chromosome arms. Crucially, although separase levels remain high upon entry into meiosis II, its inhibitor securin is substantially reduced relative to meiosis I ^16,49^. Persistent separase activity without renewed APC activation therefore creates a window of vulnerability in which premature sister chromatid separation could occur. We propose that proteasome-dependent Rec8 deprotection serves as a fail-safe mechanism to control meiosis II onset, ensuring that sister chromatid cohesion is only dissolved at the appropriate time.

A key observation underlying this model is that *pre9Δ* proteasomes retain the ability to degrade securin while being selectively impaired for shugoshin degradation. An alternative interpretation, that mutant proteasomes are equally defective toward both securin and shugoshin but cells are disproportionately sensitive to residual Sgo1 is less compelling. Were securin degradation similarly compromised, one would expect a pronounced delay in mitotic growth; however, proteasome mutants analyzed here show no such defects (**Fig. 2**), arguing against global impairment and in favor of substrate-selective degradation preference.

What accounts for this substrate selectivity? Several non-mutually exclusive mechanisms may explain why the *pre9Δ* proteasome preferentially degrades securin over shugoshin. First, the two substrates may differ in their extent, architecture, or stability of ubiquitin modifications ^50^. Polyubiquitination of securin has been experimentally demonstrated, whereas comparable evidence for yeast shugoshin or any of its orthologs is lacking, possibly because ubiquitylation of shugoshin is intrinsically unstable or rapidly reversed ^50^. In this regard, RP deubiquitinases can remove ubiquitin chains without substrate degradation ^51^, and such activity may operate in a substrate-specific manner. Second, substrate accessibility is likely to influence degradation efficiency. Securin is associated with soluble separase, rendering it accessible to the abundant nucleoplasmic proteasome pool (**Fig. 8A**). Shugoshin, by contrast, is embedded within the inner kinetochore - a densely organized structure - and this structural constraint may shield it from degradation despite polyubiquitination at metaphase I, until proteasomes are recruited to kinetochores. Consistent with this possibility, proteasomes accumulate at meiosis II kinetochores (**Fig. 8B**), suggesting that spatially restricted proteolysis contributes to the precise temporal control of chromosome segregation.

The substrate preference, uncovered here for a functional proteasome variant, may not be limited to this variant but rather reflect inherent selectivity in the wild-type complex, one that could be tuned to different developmental stages. Posttranslational modifications of the RP subunit provide a plausible mechanism for modulating the proteasome’s substrate specificity and/or localization in a stage-specific manner ^52–57^. Finally, changes in proteasome subunit composition over the course of meiosis ^58^, including the prevalence under certain stress conditions of an α4-dimer variant carrying two α4 copies in place of α3/α4 ^28^, could expand the repertoire of mechanisms by which the proteasome achieves substrate selectivity during meiosis.

### Broader implications

Our findings reveal that the proteasome regulates cohesin cleavage through two separable pathways. Whereas meiosis I chromosome segregation relies primarily on securin degradation and separase activation, meiosis II additionally requires proteasome-dependent Rec8 deprotection at centromeres. This parallel mode of proteasome action identifies meiosis II as uniquely dependent on localized proteolysis and suggests new mechanisms by which proteasome dysfunction may result in aneuploidy of gametes ^59^.

## Acknowledgements

Work in the GVB laboratory was supported by NIH grant R01GM125800. Additional funding was provided by the Center for Gene Regulation in Health and Disease (GRHD) and by a Faculty Research Development award from Cleveland State University. Work in the AM laboratory was funded by Wellcome Investigator [220780] and Discovery Awards to AM [319314], funding for the Wellcome Centre for Cell Biology [203149] and a Wellcome Discovery Research Platform Award [226791]. We gratefully acknowledge the Wellcome Discovery Research Platform for Hidden Cell Biology Proteomics Core. We thank Wolfgang Zachariae, Jasvinder Ahuja, Stefan Galander, Eva Hoffmann, Doug Bishop, and Nancy Kleckner for strains, Judith Yanowitz, Orlando Arguello-Miranda, Jesus Monge-Neria, and Weronika Borek for discussion, and Islam Muheisen, Caroline Wu, and Ashley Kucia for experimental assistance.

## Supplemental Material

**Supplemental Figures S1-S5**

**Supplemental Tables S1-S3**

**Table S1**

Spore viabilities in genotypes analyzed in this work.

**Table S2**

Excel file of significantly enriched or depleted proteins in *pre9Δ spo11-yf* at t = 6.5 h.

**Table S3**

Strains used in this study.

## Materials and Methods

### Yeast strains

Strains used in this study were of the isogenic SK1 background. Genotypes including the names of gene constructs encoding relevant fusion proteins are listed in **Table S3**. Estradiol-inducible expression from the *pGAL1* promoter was achieved with a Gal4-estrogen receptor fusion controlled by the constitutive *pGPD1* promoter ^60^. Alleles *rec8-mNeonGreen* (*rec8-mNeon*) and *sgo1-V5_3_* (gifts from Stefan Galander), *sgo1*-*mNeonGreen* (*sgo1-mNeon*)^26^, and *mtw1*-*tdTomato* were previously described ^26^. Allele *V5*-*rec8-TEV287* which carries a V5 tag at its N-terminus and a TEV protease cleavage site at amino acid 287, as well as *pGAL1* promoter-driven TEV protease were gifts from Eva Hoffmann ^39^. *rec8-14D-GFP* and *rec8-GFP* were gifts from Wolfgang Zachariae ^15^. Double and triple mutant strains were constructed using standard mating and sporulation procedures followed by mating of haploid spore clones carrying the desired allele combinations. All strains were verified by PCR-based genotyping and/or fluorescence marker analysis where appropriate.

### Meiotic time course

Unless noted, nuclear divisions, chromosome segregation, immunocytological preparations, protein preparations, and spore viability, were performed using synchronized meiotic cultures. Meiotic time courses were established by pre-growth in YPA (1% yeast extract, 2% peptone, 1% potassium acetate) for 13.5 h at 30°C, followed by transfer to meiosis medium (a.k.a. sporulation medium; 0.5% potassium acetate, 0.02% raffinose, and 3 drops antifoam per liter) ^61^. For *rpt6*-HA_3_ strains, all pre-meiotic growth steps, including propagation on YPG and YPD plates and subsequent growth in YPA, were performed at 33°C. Following transfer to meiosis medium, all strains including *rpt6*-HA_3_ were incubated at 23°C. Pre-meiotic cultures were transferred to meiosis medium pre-equilibrated at 23°C and incubated for the indicated times. *PDR5* encodes a multidrug transporter responsible for cellular export ^62^. For investigating effects of MG132, a single premeiotic *pdr5Δ* culture was incubated in meiosis medium at 23°C until t = 7 h, followed by a split into two equal parts, with one part receiving the indicated amount of MG132 (Calbiochem, #474791) dissolved in DMSO (Fisher, #BP231-1), or DMSO only (mock).

#### Induction of NDT80

High levels of *NDT80* expression were achieved in strains carrying *pGAL1-ndt80* by adding 100 μM β-estradiol (Sigma, #E8875) at t = 7.5 h, a time when the majority of wild-type cells has reached the prophase I arrest point at 23°C.

#### Induction of TEV protease

High levels of TEV protease expression was achieved in strains carrying *pGAL1-NLS-myc_9_-TEV*-protease by adding 100 μM β-estradiol at t = 9.5 h when *pre9Δ SPO11* cultures exhibit peak levels of binucleate cells at 23°C.

### Tetrad dissection and spore viability analysis

Spore viabilities were determined from the same synchronized meiotic cultures used for the corresponding cytological analyses. Following spore formation, cells were treated with zymolyase (0.5 mg/ml) to digest the ascus wall and tetrads were dissected on YPD plates using a micromanipulator (Zeiss). Dissected spores were incubated at 30°C for 2 to 3 days to allow spore colony formation. Tetrad viability was determined by scoring the number of viable colonies arising from each tetrad; overall viability was calculated as the percentage of spore colonies out of the number of spores dissected.

### Western blot analysis in yeast extracts

Primary antibodies were mouse anti-V5 (Biorad #MCA1360GA) (1:1,000), mouse anti-PGK1 (Abcam, #ab113687) (1:2,000), mouse anti-myc tag antibody (9E10) (Abcam, #ab32) (1:500) and Rabbit Anti-α tubulin antibody (EPR13799; Abcam#184970) (1:10000). Secondary antibodies used were goat anti-mouse IgG H&L-HRP (Abcam, #ab205719) (1:2,000) and Goat anti-Rabbit IgG, HRP conjugate (Merck#12-348) (1:2000). For western immunoblotting, samples were processed as described ^63^. Samples were separated on precast 4-20% SDS-PAGE gels (BioRad #4561096) in SDS running buffer and blotted to PVDF membranes (0.45 μM, Millipore #IPFL00010) in transfer buffer in a Bio-Rad Mini Trans-Blot system. Membranes were blocked in 5% dry milk powder in PBS with 0.05% Tween20 (PBST) for at least 1 h before incubating with primary antibody in 5% dry milk powder/PBST overnight at 4°C. Membranes were washed in PBST three times for 15 min, incubated with secondary antibody in 5% dry milk powder/PBST for 1 h at room temperature, and washed in PBST three times. HRP-conjugated antibodies were detected with a chemiluminescent substrate (Thermoscientific #34577) using a LI-COR imaging system, and images were quantitated using ImageJ (National Institutes of Health).

### Mass spectrometry analysis

#### Meiotic culturing

Strains were patched from -80°C to YPG plates (1% yeast extract, 2% Bacto^TM^ peptone, 2.5% glycerol, and 2% agar) and incubated at 30°C for 24 h. Next, two 12.5 ml YPDA (1% yeast extract, 2% Bacto^TM^ peptone, 2% glucose, 0.3 mM adenine) cultures were inoculated from YPG plates and grown at 30°C with shaking at 250 rpm for 24 h. Next, two 50 ml BYTA (1% yeast extract, 2% Bacto^TM^ tryptone, 1% potassium acetate, 50 mM potassium phthalate) cultures were started at OD_600_∼0.3 and shaken at 250 rpm at 30°C overnight (∼16 h). Cells were harvested, washed twice with 25 ml water, resuspended to make six 25 ml SPO (0.3% potassium acetate) cultures (three per strain) to an OD_600_∼2.0. Nuclear divisions were monitored at the indicated time points. At t = 6.5 h, 10 ml of culture was collected for TCA protein extracts. The culture for TCA extract preparation was spun at 3.000 rpm for 2 min and resuspended in 5 ml 5% TCA on ice. Then the samples were spun again at 3,000 rpm for 2 min, transferred to a Fastprep^TM^ tube, spun at 13,200 rpm for 1 min and the cell pellets drop frozen in liquid nitrogen and stored at -80°C.

#### Sample preparation

Frozen cell pellets were washed once in 0.5 ml ice-cold acetone and spun for 10 min at 13,200 rpm in a microcentrifuge. Acetone was carefully poured and pipetted off pellets in a hood and allowed to dry for 10 min. Cell pellets were resuspended in 0.2 ml of urea lysis buffer (8 M urea, 2 mM Pefabloc, 2 mM b-glycerophosphate, 1 mM Na pyrophosphate, 5 mM NaF, 0.8 mM sodium orthovanadate, 10 μg/ul each of ‘CLAAPE’ (chymostatin, leupeptin, antipain, aprotinin, pepstatin A, E-64 protease inhibitor), 0.2 μM microcystin, and 1x Roche complete protease inhibitor cocktail. Silica beads were added, and bead-beating lysis was performed with a Fastprep machine (MP Biomedicals) with 4 rounds of lysis, 2 min on ice in between. Lysates were transferred to a new tube by poking a hole with a red-hot needle and spinning for 20 sec into a new tube. Lysates were spun in a 4°C microcentrifuge at 7,000 rpm for 15 min and supernatant collected. Protein concentrations were measured by BCA assay and 400 μg per sample was transferred to a new tube. Samples were reduced with 5 mM DTT for 25 min at 37°C, then treated with 10 mM IAA in the dark at room temperature, followed by 15 mM DTT in the dark for 15 min at room temperature. Protein was precipitated with 500 ul of ice-cold acetone at -20°C overnight. The next day, samples were spun at 8,000 rpm for 10 min at 4°C and dried in a fume hood. Samples were resuspended in 100 mM fresh TEAB and 5 μg trypsin added, incubated at 37°C for 4 h and then another 5 μg trypsin added and mixed by vortexing before incubating overnight at 37°C. On the next day, peptides were collected by spinning in a microcentrifuge at 4,000 rpm for 10 min and collecting the supernatant. Trypsin-digested supernatants were brought to a volume of 100 μl in 100 mM TEAB. Each 400 μg peptide sample was mixed with 0.8 mg TMT label resuspended in 41 μl ACN and incubated at 25°C for 1 h, shaking at 400 rpm. Labelling reactions were stopped by adding 50 mM Tris pH 8.0 and incubated at 25°C for 15 min, shaking at 400 rpm. Next, the six samples were combined in 1 tube and acidified with 100% formic acid until pH <3. Next, samples were dried in a vacuum centrifuge to completion and then stored at -80°C. Peptide sample was subsequently thawed and resuspended in 0.2% formic acid and desalted using a 500 mg C18 SepPak column (Waters). 4% of the sample was reserved and loaded onto a C18 stage tip for preliminary analysis and the remaining sample was fractionated by RP-HPLC into 12 fractions. 12 fractions were loaded onto C18 stage tips and analysed on an Orbitrap Fusion^TM^ Lumos^TM^ MS (WCB Proteomics, University of Edinburgh).

#### Data analysis

All data analysis was performed using the proteinGroups.txt from MaxQuant using R (Bioconductor) within RStudio environment. R analysis done by using DEP (Differential Enrichment Analysis of Proteomics Data) package. Differentially enriched and depleted proteins identified were subjected to Gene Ontology (GO) enrichment analysis using the R package gprofiler2. Significantly enriched GO terms were retrieved from the g:Profiler database, and results were visualized by plotting GO terms against their enrichment significance, expressed as the negative logarithm of the adjusted p-value (-log10 p-value). Separate analyses were performed for proteins enriched and depleted in the mutant relative to wild type.

### Microscopy

Unless otherwise indicated, images were acquired using a Nikon Eclipse Ti2 microscope equipped with a 100×/1.45 NA Plan Apo oil objective, an Abberior STEDYCON imaging system, and a motorized stage. Z-stack images were acquired using identical imaging settings within each experiment and processed as maximum-intensity projections for analysis.

#### Fluorescence microscopy analysis

Cell aliquots from liquid meiotic cultures were collected at appropriate time points, fixed as described, and stained with 1 μg/mL 4′,6′-diamidino-2′-phenylindole (DAPI) at room temperature. Meiotic divisions were scored by counting the number of DAPI-staining nuclei per cell using a Nikon Eclipse Ti2 fluorescence microscope.

Segregation of *cen5*-GFP or *cen3*-GFP-tagged centromeres was determined in binucleate or tetranucleate yeast cells by mixing one volume of cells with one volume of fixative solution [3% (w/v) paraformaldehyde; 3.4% (w/v) sucrose], spreading the mixture with a glass rod on a microscope slide, staining slides in TBS containing 1 μg/ml DAPI and photographing cells soon thereafter, using a widefield microscopy fluorescence imaging system (Nikon Eclipse Ti2). For binucleate cells, aliquots from t = 6 h, 7.5 h and 9 h were analysed, and for tetranucleate cells the t=24 h time point was used.

#### Imaging of fixated intact cells

Cells were induced to undergo synchronous meiosis as described in the “Meiotic time course” section. At the indicated time points, aliquots were collected, fixed, and imaged as described under “Microscopy”. Mtw1-tdTomato was used as a kinetochore marker to define meiotic stages. Fluorescence intensities were quantified using STEDYCON image analysis software (Abberior) by measuring the mean signal intensity within a 1-μm-diameter circular region of interest (ROI) centred on individual kinetochore foci. Local background fluorescence was subtracted from each measurement. Background-corrected fluorescence intensities were averaged for each meiotic stage.

#### Live-cell imaging

Cells were prepared for synchronous meiosis as described in the “Meiotic time course” section. After 5 h in 0.5% potassium acetate meiosis medium (above), cells were transferred to concanavalin A-coated 8-well glass-bottom dishes (Ibidi^TM^), allowed to adhere for 20 min, followed by addition of fresh 2% SPM. Following a 20-min equilibration period on the microscope stage, z-stack images were acquired every 10 min for 2 to 2.5 h at room temperature using identical imaging settings throughout each experiment. Maximum-intensity projections were generated for analysis. Fluorescence intensity measurements were performed in NIS-Elements General Analysis software (Nikon). Total Rec8-mNeon fluorescence intensity was measured within a region of interest (ROI) encompassing the entire cell. Local background fluorescence was determined from an adjacent region of identical size and subtracted from the ROI measurement.

#### Surface chromosome squashes and immunofluorescence microscopy

Surface chromosome squashes were prepared similarly as surface chromosomes spreads described previously ^61^, except that nuclei were squashed by gentle compression under a coverslip rather than by dispersion with a glass rod. Pup2-GFP and Mtw1-tdT were detected using goat anti-GFP (Rockland, 600-101-215; 1:200) and rabbit anti-mCherry (Abcam, ab167453; 1:200) primary antibodies, respectively, followed by donkey anti-goat Alexa Fluor 488 (Invitrogen A-11055; 1:300) and donkey anti-rabbit Alexa Fluor 594 (Abcam, ab150064; 1:500) secondary antibodies. Images were acquired as described under “Microscopy”. Images were acquired as single optical sections. Proteasome–kinetochore association was determined by assigning Pup2-GFP foci and Mtw1-tdTomato signals to 30° angular sectors within DAPI-stained nuclei, and frequencies in each sector were determined thereafter.

### Statistical analysis

Statistical analyses were performed using GraphPad Prism (version 10; GraphPad Software). Data are presented as mean ± SD, mean ± SEM, or box-and-whisker plots, as indicated in the corresponding figure legends. The statistical tests used for each dataset are specified in the respective figure legends. Biological replicates (N) and sample sizes (n) are indicated in the figures or figure legends. A *P* value of < 0.05 was considered statistically significant.

---

**Fig. S1.**
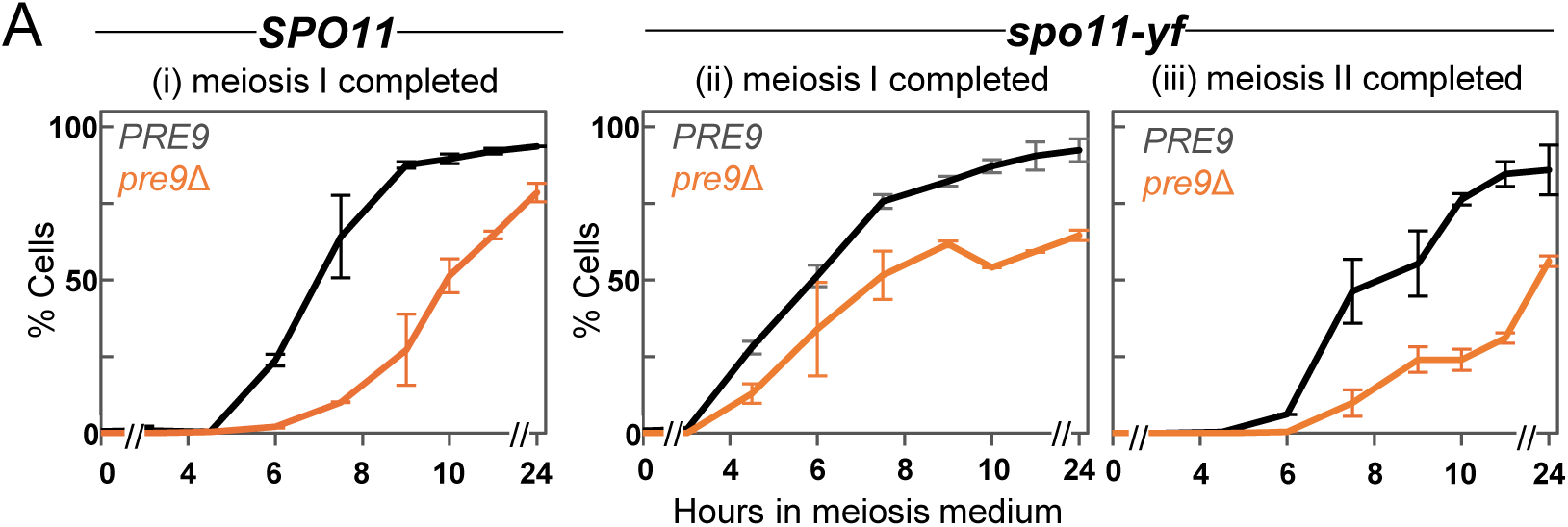
Effects of *pre9Δ* (i) in *SPO11* on meiosis I nuclear divisions; and in *spo11-yf* on (ii) meiosis I nuclear divisions, and (iii) meiosis II nuclear divisions.

**Fig. S2.**
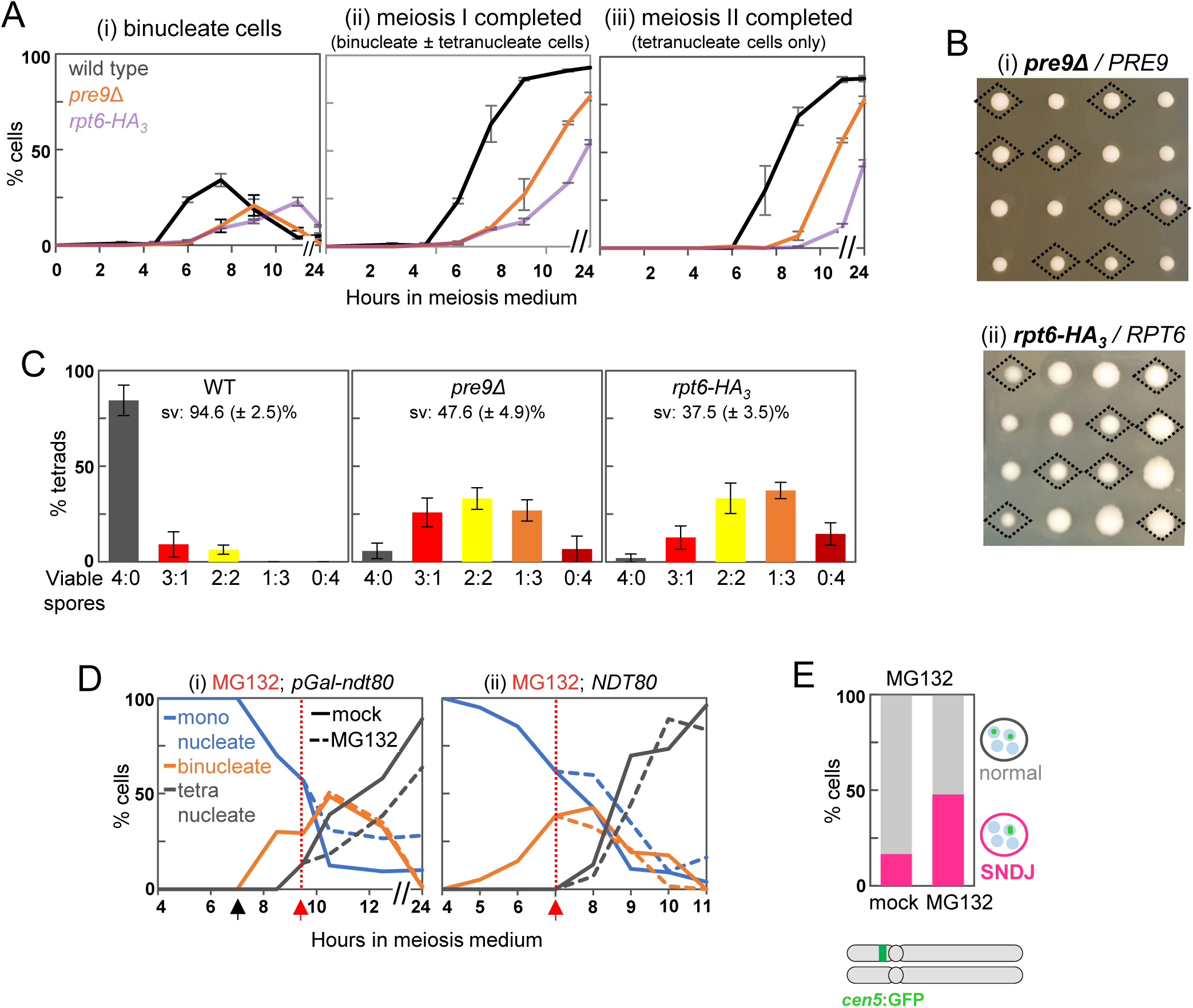
Meiosis II sister chromatid disjunction is distinctly sensitive to impaired proteasome function. (A) Progression through meiosis in wild type, pre9Δ and rpt6-HA3. (i) Steady-state levels of binucleate cells, (ii) cumulative levels of cells that had at least completed meiosis I, (iii) meiosis II completion. Cultures were incubated at 23°C (n = 2). (B) Mitotic growth of colonies derived from individual haploid spores from heterozygous strains (i) pre9Δ::KanMX4/ PRE9 and (ii) rpt6-HA3 ::HygroMX4/ RPT6. Each row shows the four products of a single meiosis which yielded two spore colonies carrying the (i) KanMX4- or (ii) HygroMX4-marked proteasome allele (diamonds), and two spores carrying the respective unmarked wild-type allele. Growth plates were incubated at (i) 30°C or (ii) 34°C. (C) Tetrad viabilities in wild type (N = 6; n = 300), pre9Δ (N = 5; n = 272), and rpt6-HA3 (N = 2; n = 153) in cultures analyzed for GFP segregation in **Fig. 2**. Tetrads formed following incubation at 23°C in liquid meiosis medium were incubated and assessed for viability. Spore viabilities are averages for the indicated numbers of meiotic cultures. Error bars indicate ranges for each tetrad viability class. Ratios indicate viable: inviable spores. (D) Effects of MG132 treatment on meiotic divisions in cells carrying (i) inducible NDT80 (pGal1-ndt80) or (ii) wild-type NDT80. MG132 in DMSO or DMSO was added when binucleate cells had reached maximum levels (i) at t = 9.5 h in pGal1-ndt80 following NDT80 induction with β-estradiol at t = 7 h; and (ii) at t = 7 h in NDT80. Black arrow indicates NDT80 induction, red arrows indicate addition of MG132 at (i) 60 μM or (ii) 20 μM. All strains are also pdr5Δ/”. Cultures were incubated at 23°C. (E) Frequencies of sister chromatid non-disjunction following completion of meiosis II as inferred from distribution of heterozygous GFP signals associated with cen5 in mock- or MG132-treated pdr5Δ/” strains (mock, n = 102; MG132, n = 109).

**Fig. S3.**
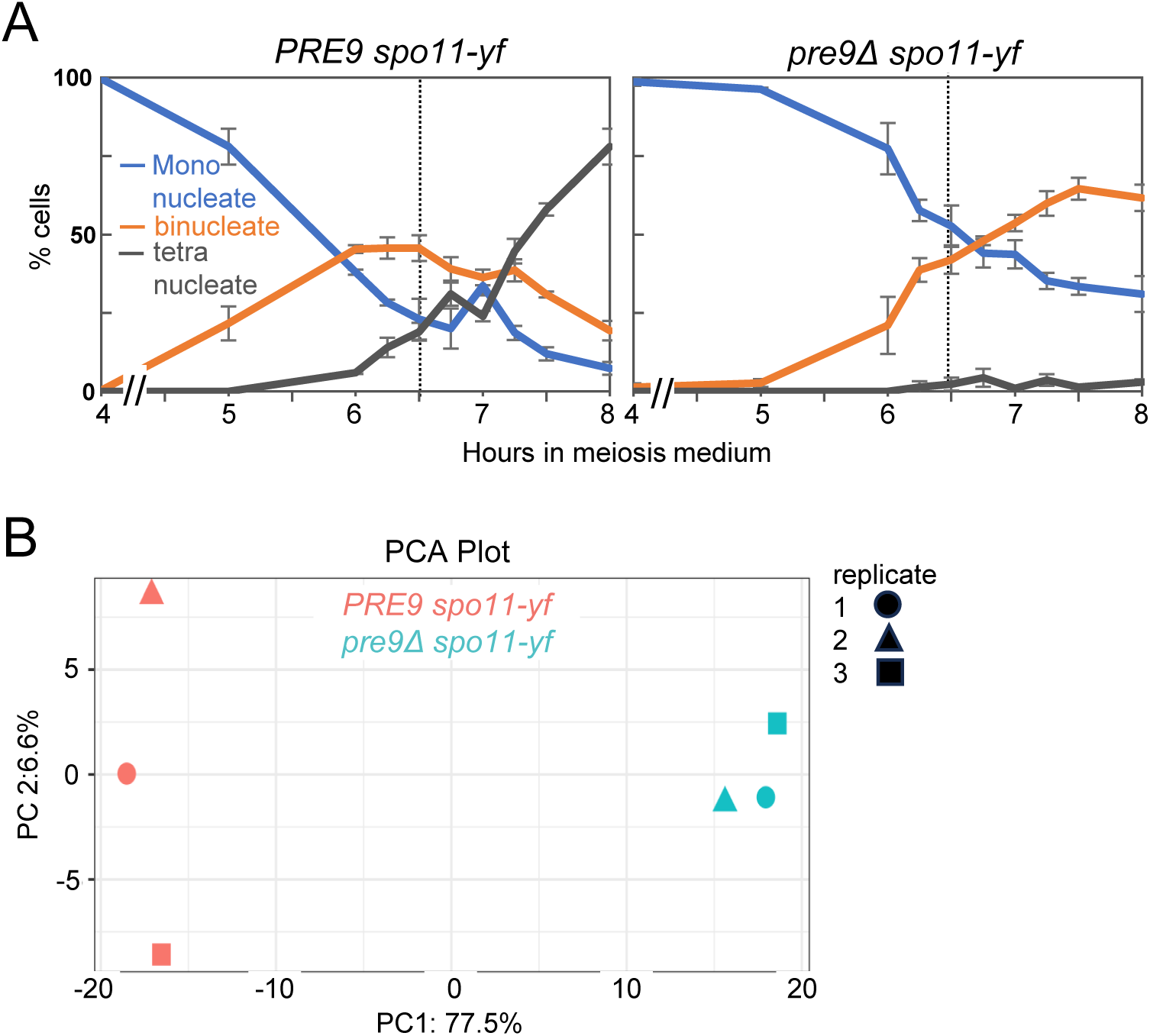
Mass spectrometry analysis of proteome in *pre9Δ* proteasome mutant. (A) Meiotic divisions in cultures of PRE9 and pre9Δ in the spo11-yf background at 30℃, sampled at t = 6.5 h (see dotted line) for mass spectroscopy. n = 3, error bars indicate range. (B) Principal component analysis (PCA) of top 597 variable proteins. PC1 captures variation between PRE9 spo11-yf and pre9Δ spo11-yf samples on global protein expression, PC2 captures variation between biological replicates.

**Fig. S4.**
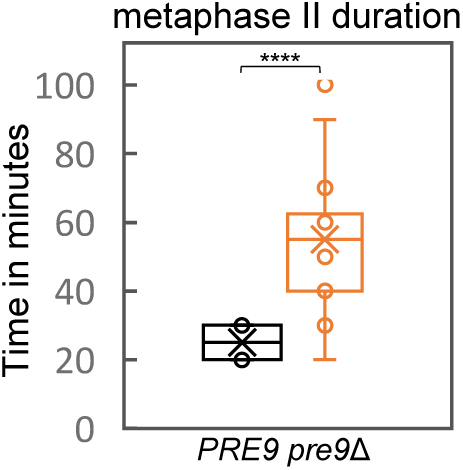
Metaphase II duration in live-cell imaging. Duration of the metaphase II stage, measured by live-cell imaging from appearance of the 2*2Kt stage to the time when individual kinetochores within a pair have separated by more than 1 μm (4Kt; n = 18). Box-and-whisker plots show the median (center line), the interquartile range (box), whiskers extending to 1.5 × interquartile range, and individual outliers shown as points. Significance was determined by unpaired t test with Welch’s correction; ****, p < 0.0001.

**Fig. S5.**
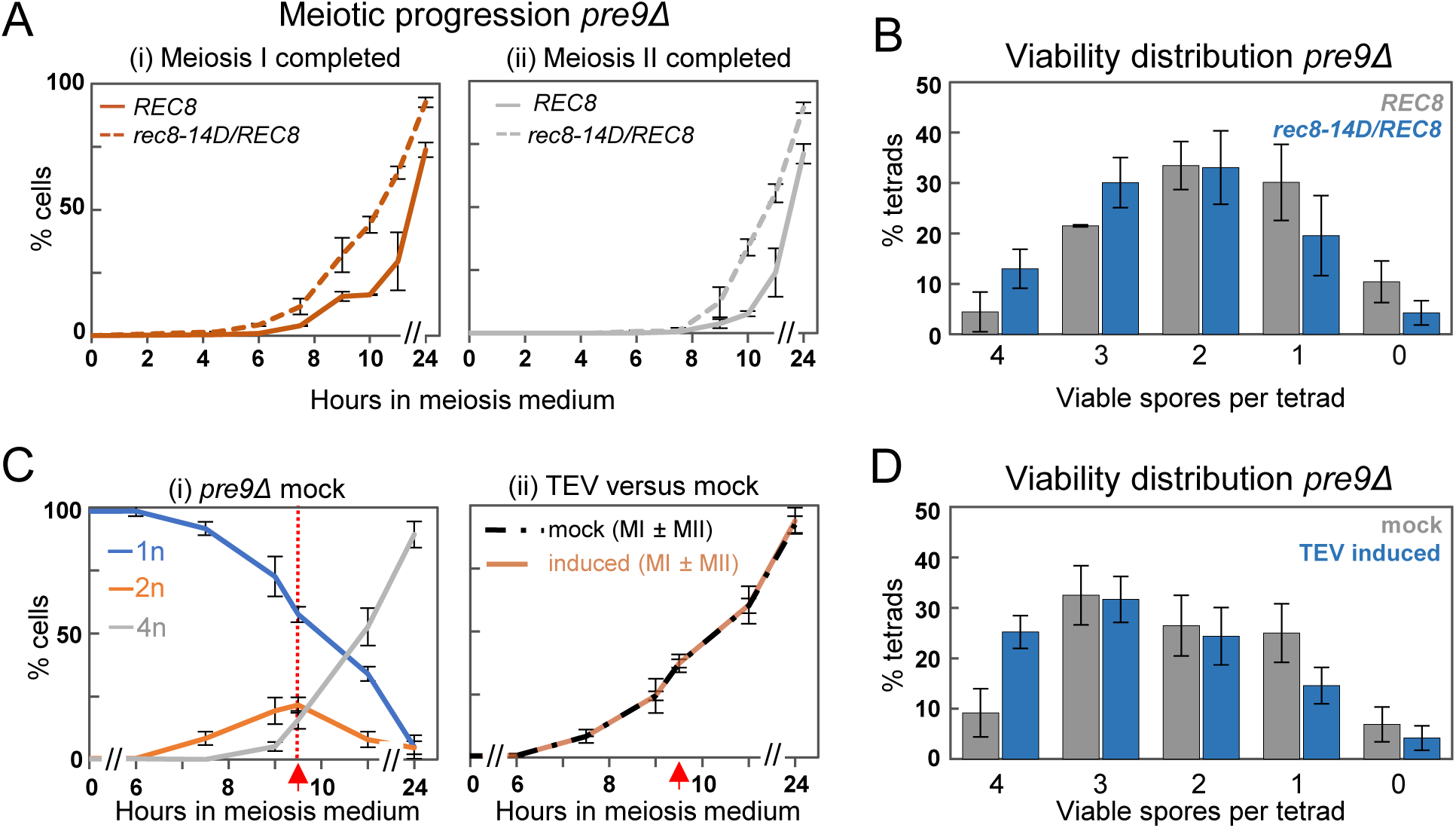
Effects of phosphomimetic Rec8 or induced Rec8-cleavage on meiotic progression and spore viability. (A) Average meiotic divisions in pre9Δ strains heterozygous for rec8-GFP/REC8 and rec8-14D-GFP/REC8, respectively. (i) Cumulative completion of meiosis I; (ii) completion of meiosis II (n = 3). (B) Viability patterns in tetrads derived from meiotic cultures shown in **Fig. S5A-ii.** (C) Average meiotic divisions in pre9Δ strains carrying TEV protease under the control of the pGal1 promoter. Red arrows indicate time of addition of TEV-inducer β-estradiol (i) Proportions of 1, 2, or 4 nuclei in mock-induced cultures. (ii) Meiosis I completion in induced compared to mock cultures (n = 3). (D) Viability patterns in tetrads derived from meiotic cultures shown in **Fig. S5C-ii**

**Table S1.**

|  | Genotype | Strain | Tetrad Dissected | Viable Spores | Total Spores | Spore Viability |
| --- | --- | --- | --- | --- | --- | --- |
| 1. | *PRE9*/ “ | AMY10 | 81 | 309 | 324 | 95.37% |
| 2. | *PRE9*/ “ | AMY11 | 20 | 71 | 80 | 88.75% |
| 3. | *PRE9*/ “ | AMY939 | 200 | 759 | 800 | 94.87% |
| 4. | *pre9Δ*/ “ | AMY9 | 132 | 242 | 528 | 45.83% |
| 5. | *pre9Δ*/ “ | AMY12 | 20 | 42 | 80 | 52.50% |
| 6. | *pre9Δ*/ “ | AMY396 | 120 | 213 | 480 | 44.37% |
| 7. | *rpt6-HA<sub>3</sub>*/ “ | AMY941 | 153 | 229 | 612 | 37.53% |
| 8. | *V5-rec8*/ “<br>(TEV Uninduced) | AMY669 | 40 | 140 | 160 | 87.5% |
| 9. | *V5-rec8*/ “<br>(TEV induced) | AMY669 | 40 | 149 | 160 | 93.12% |
| 10. | *V5-rec8*/ “, *pre9Δ*/ “<br>(TEV Uninduced) | AMY689 | 200 | 426 | 800 | 53.25% |
| 11. | *V5-rec8*/ “, *pre9Δ*/ “<br>(TEV induced) | AMY689 | 200 | 519 | 800 | 64.87% |
| 12. | *rec8-mNG*/ “ | AMY839 | 10 | 39 | 40 | 97.5% |
| 13. | *rec8-14D-GFP/rec8-14D-GFP* | AMY1216 | 40 | 2 | 160 | 0.125% |
| 14. | *rec8-GFP/REC8* | AMY1442 | 20 | 78 | 80 | 97.5% |
| 15. | *rec8-14D-GFP/REC8* | AMY1446 | 20 | 72 | 80 | 90% |
| 16. | *rec8-GFP/REC8*, *pre9Δ*/ “ | AMY1297 | 240 | 426 | 960 | 44.37% |
| 17. | *rec8-14D-GFP/REC8*,<br>*pre9Δ*/ “ | AMY1291 | 320 | 733 | 1280 | 57.26% |

**Table S2.**
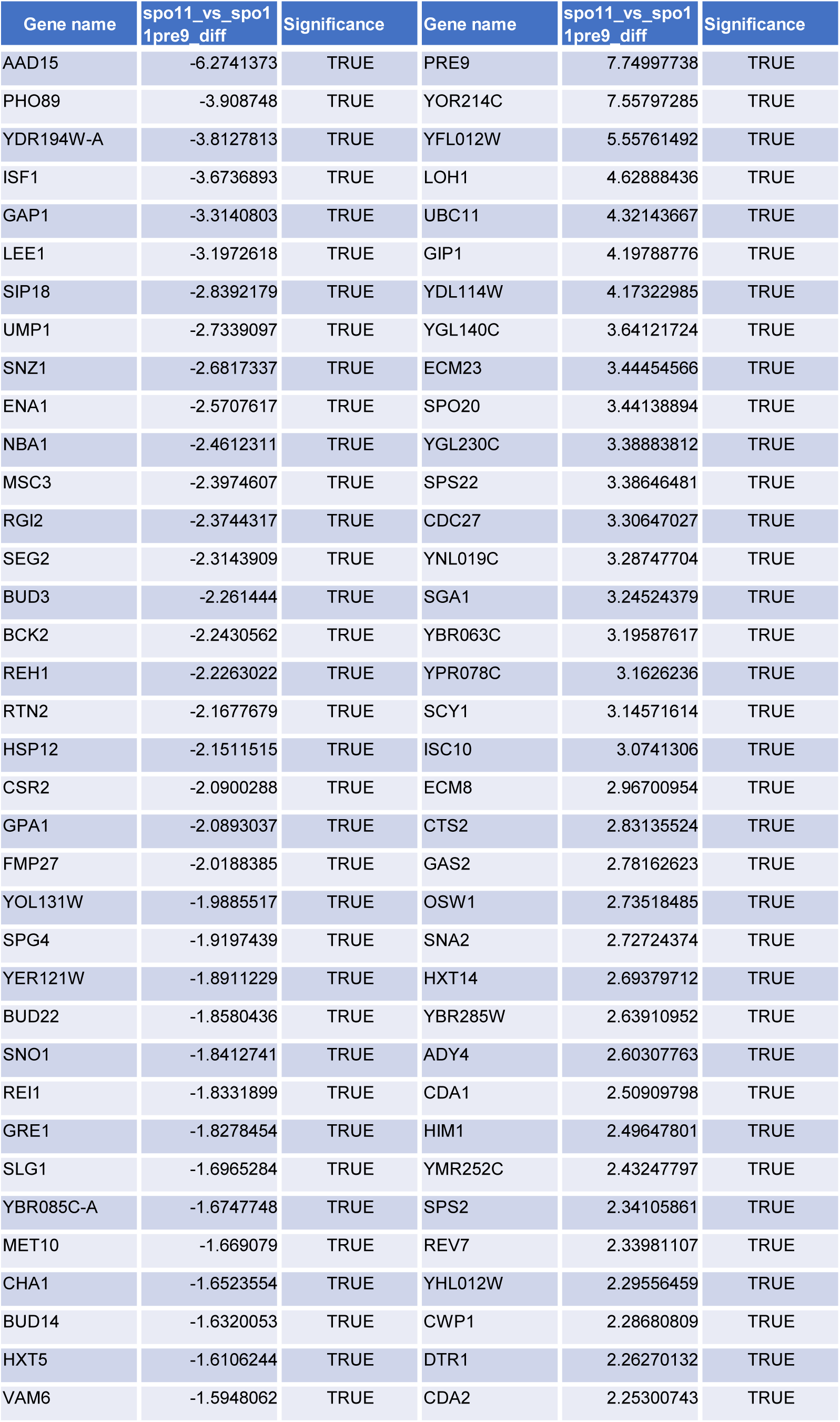
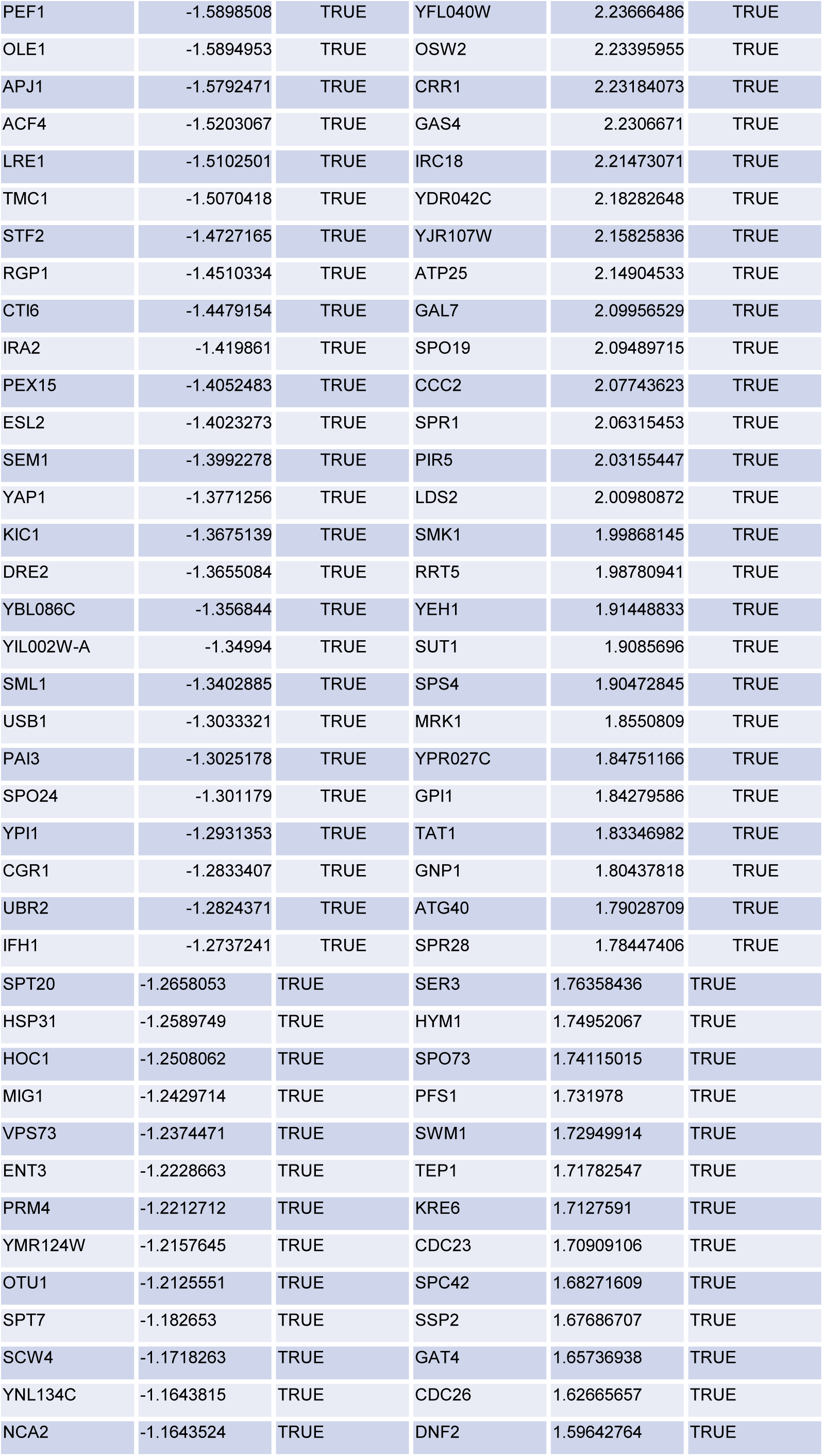
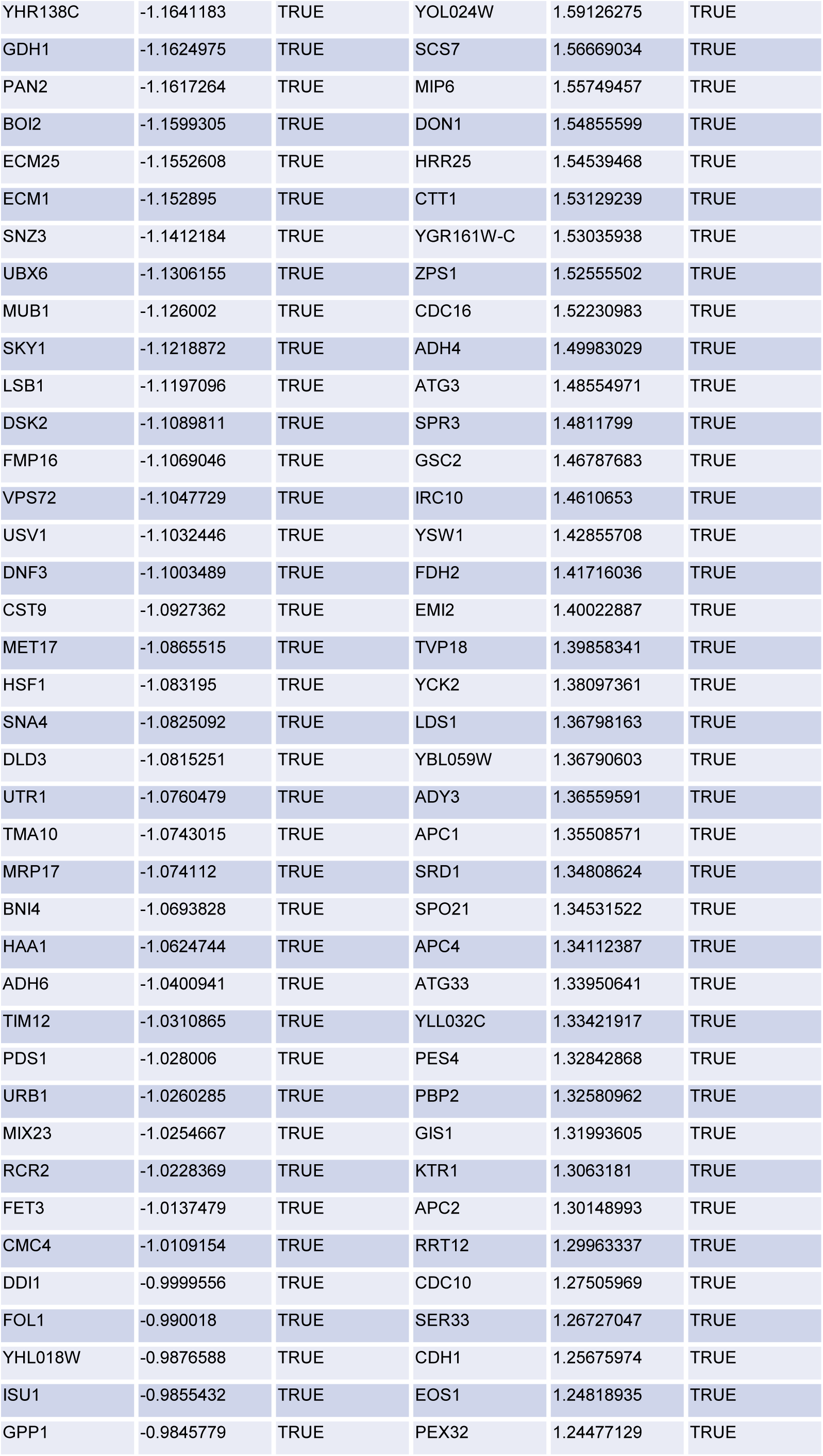
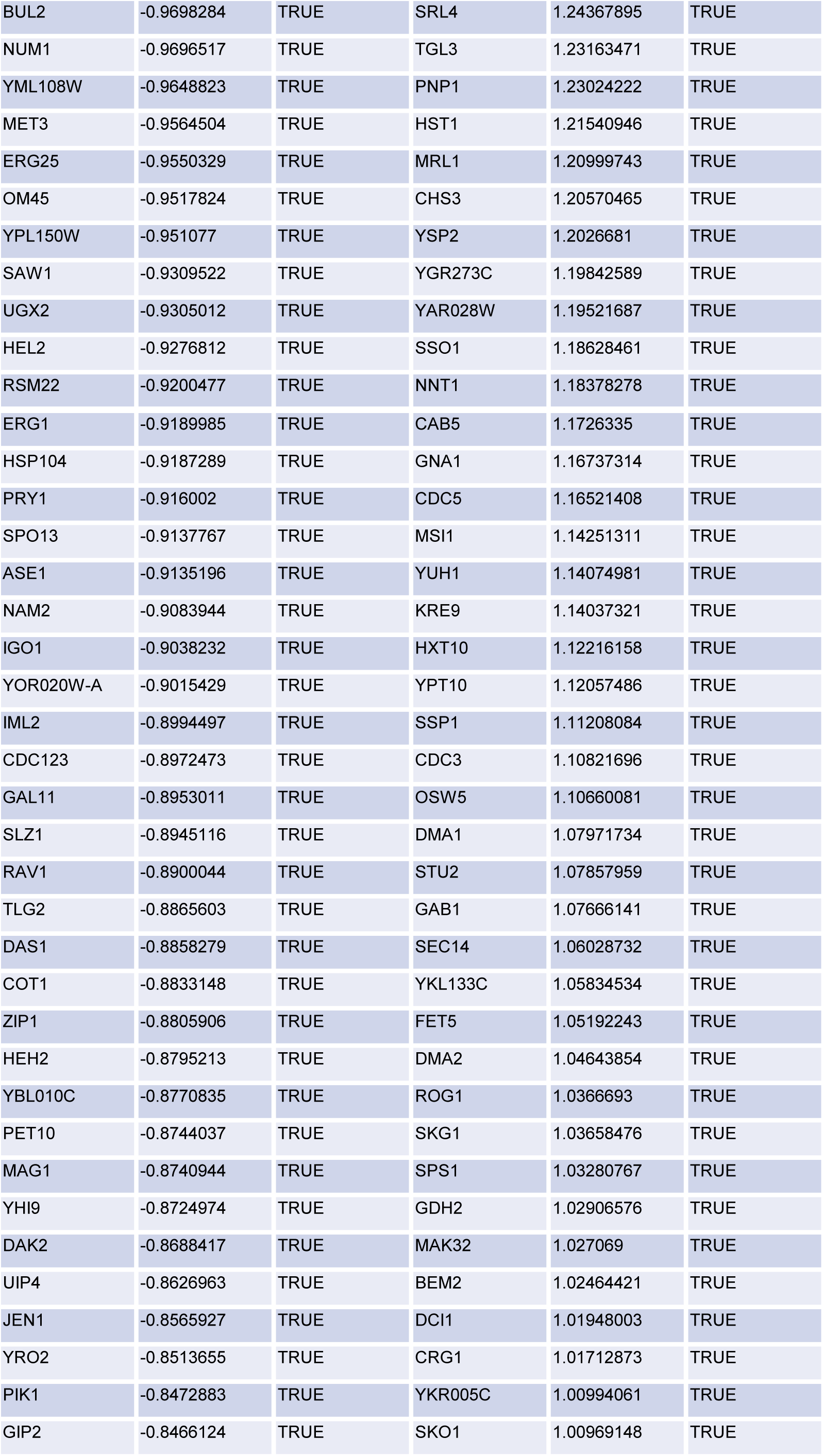
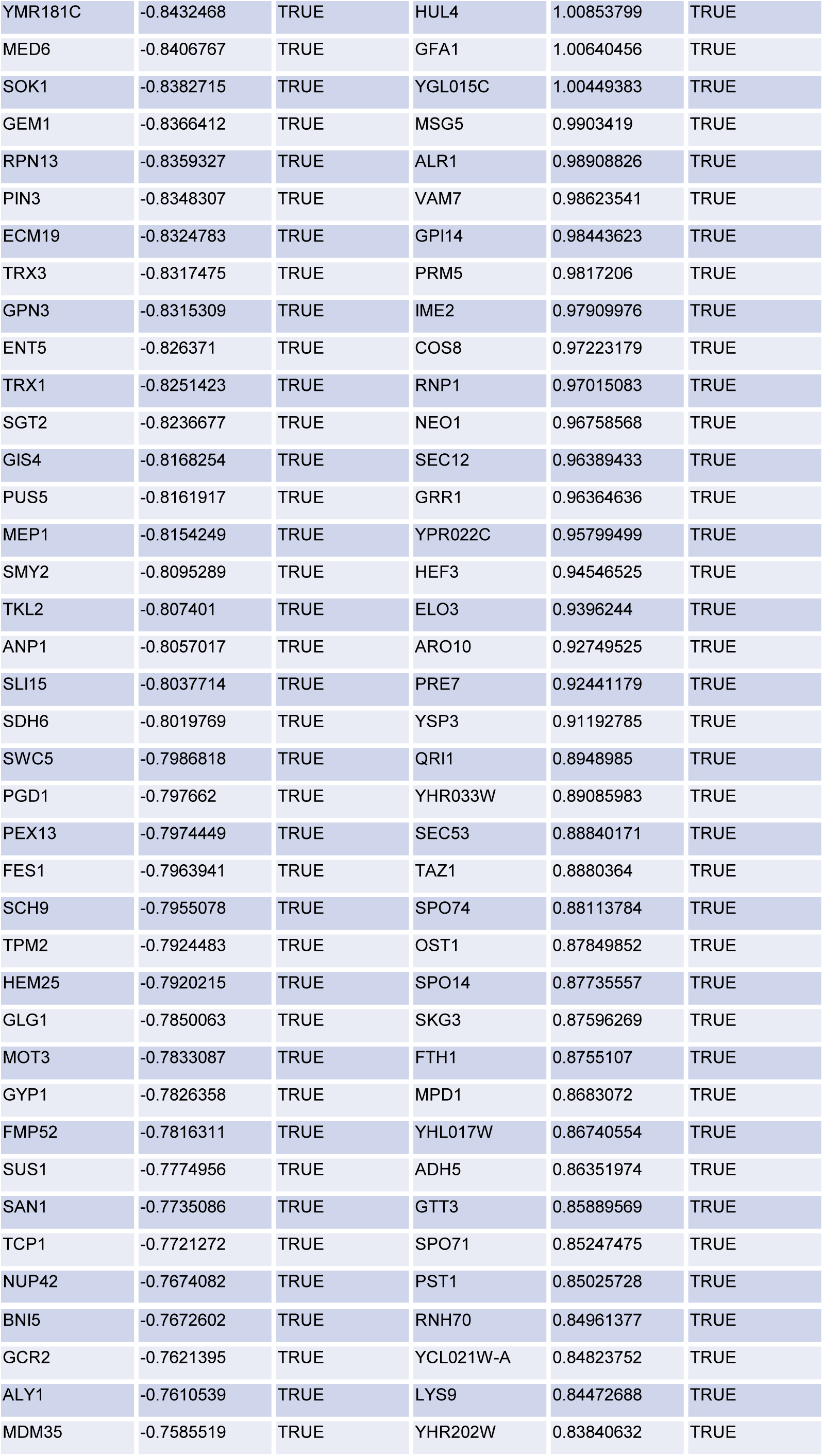
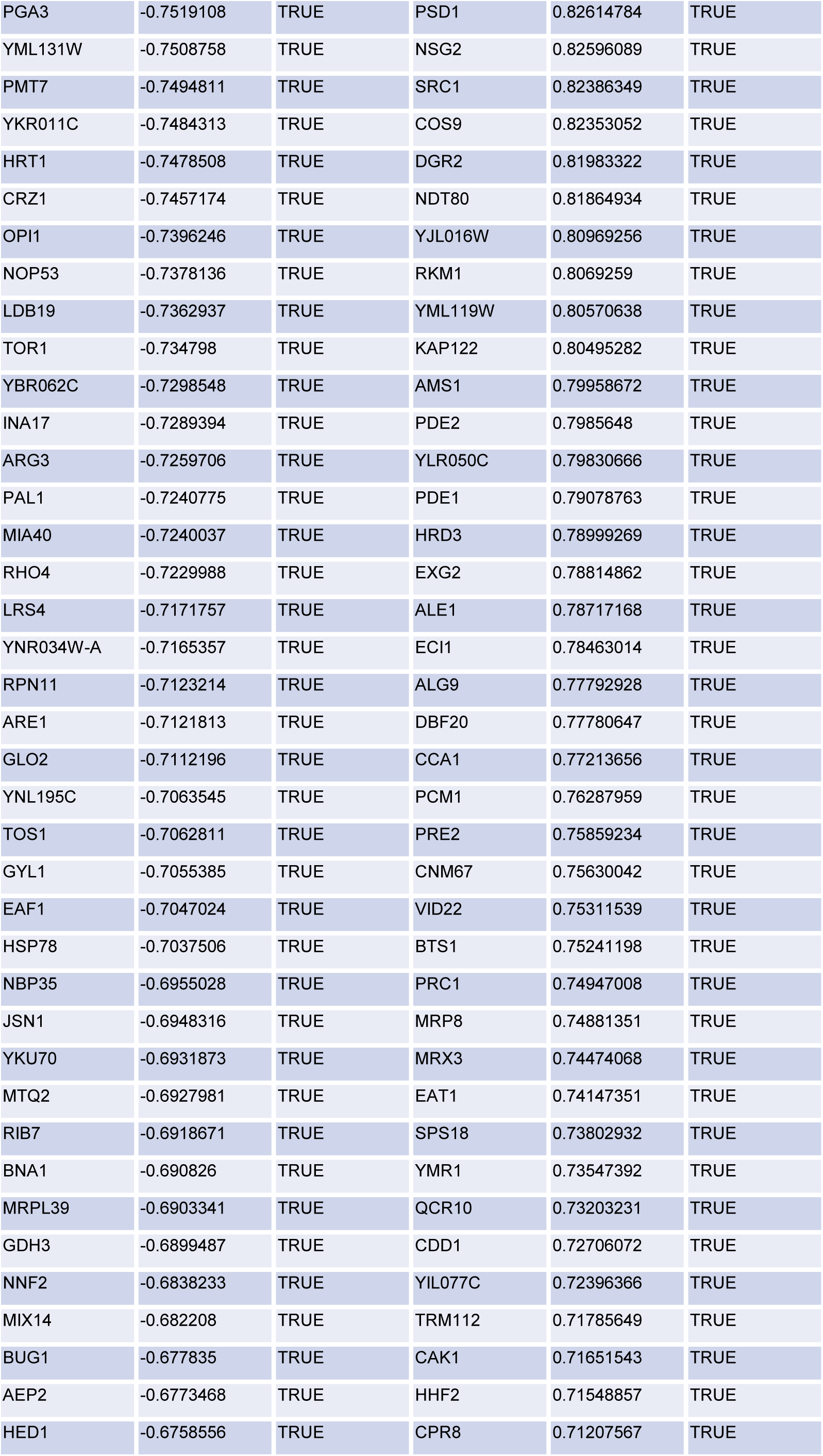
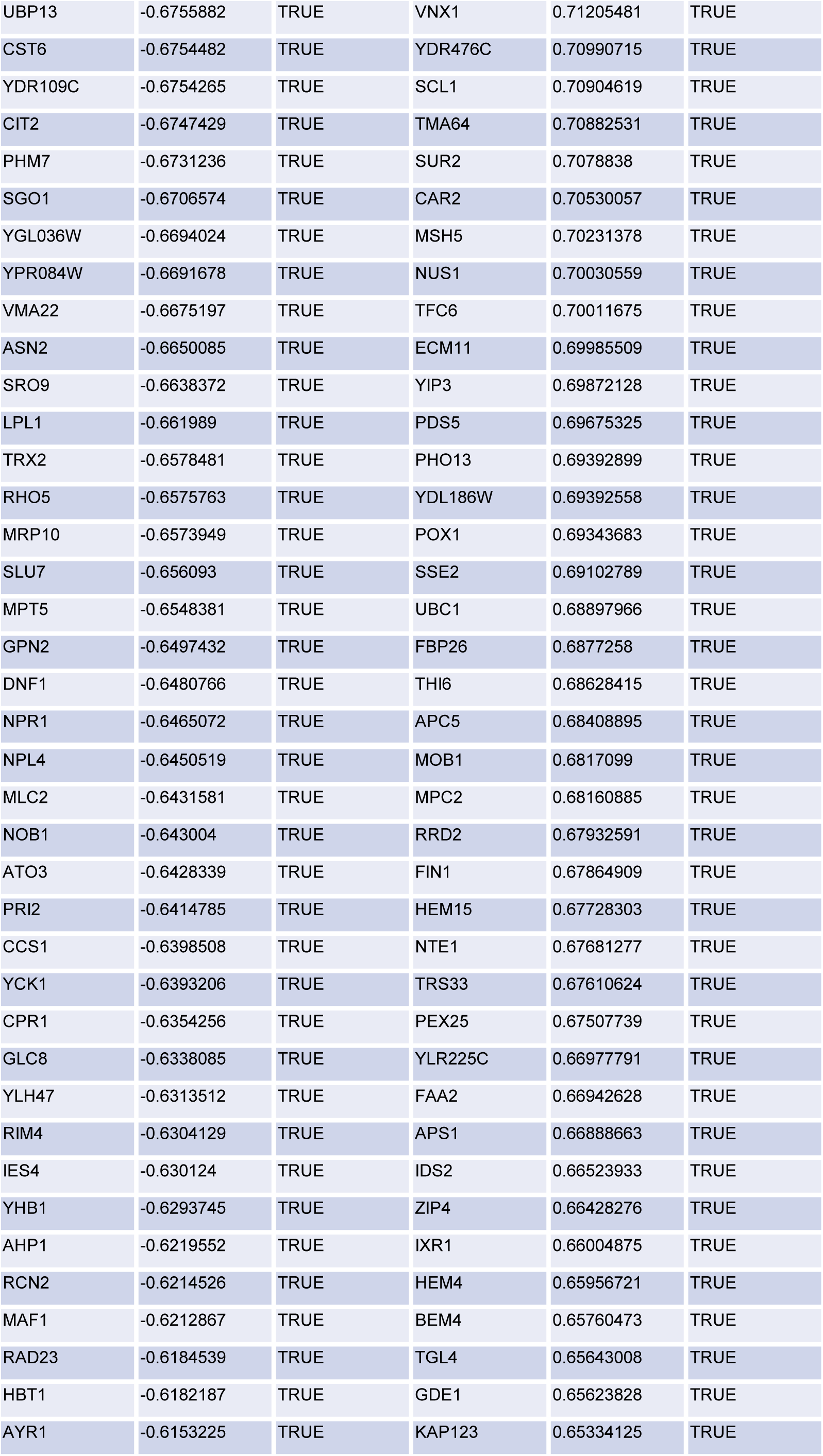
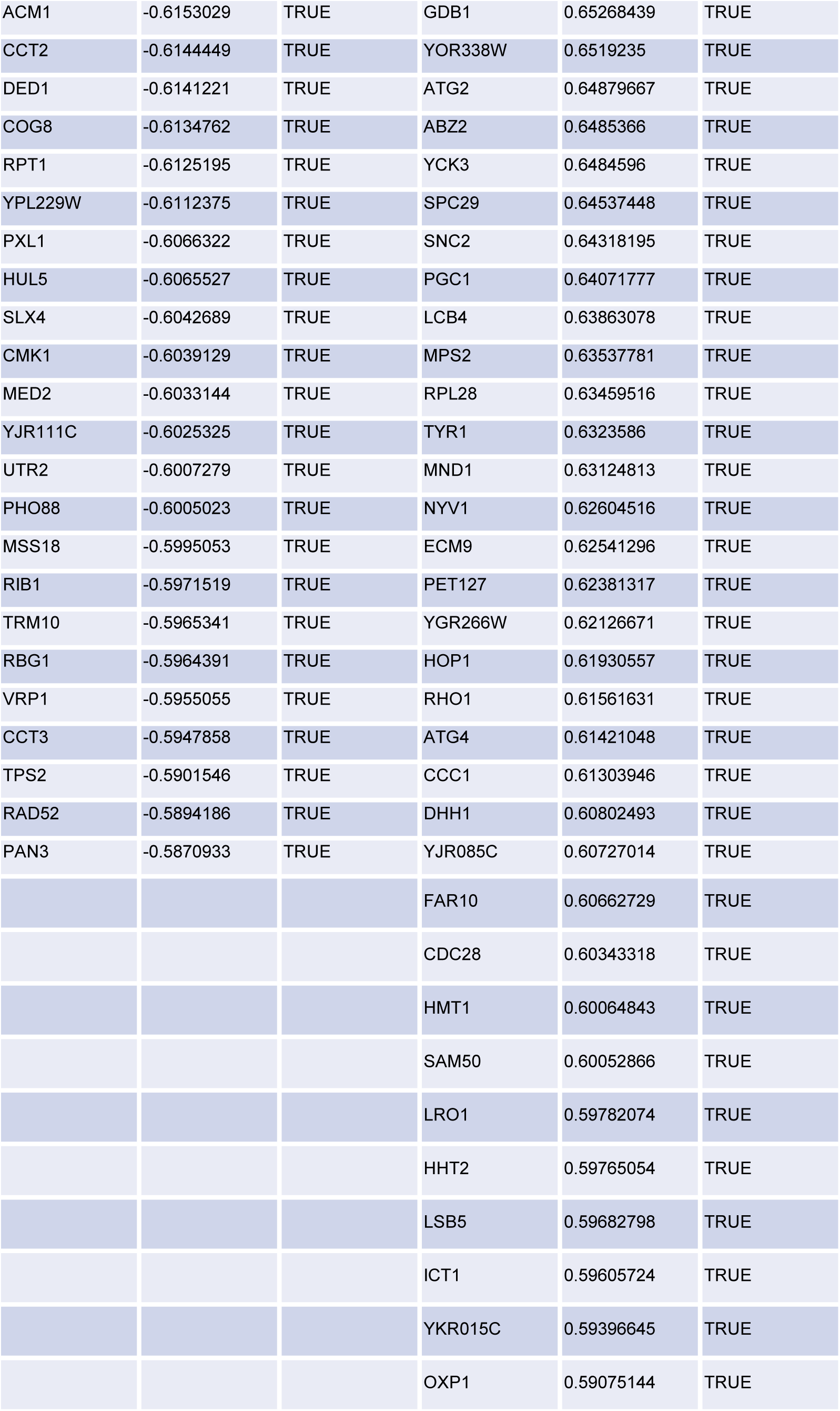

**Table S3.**
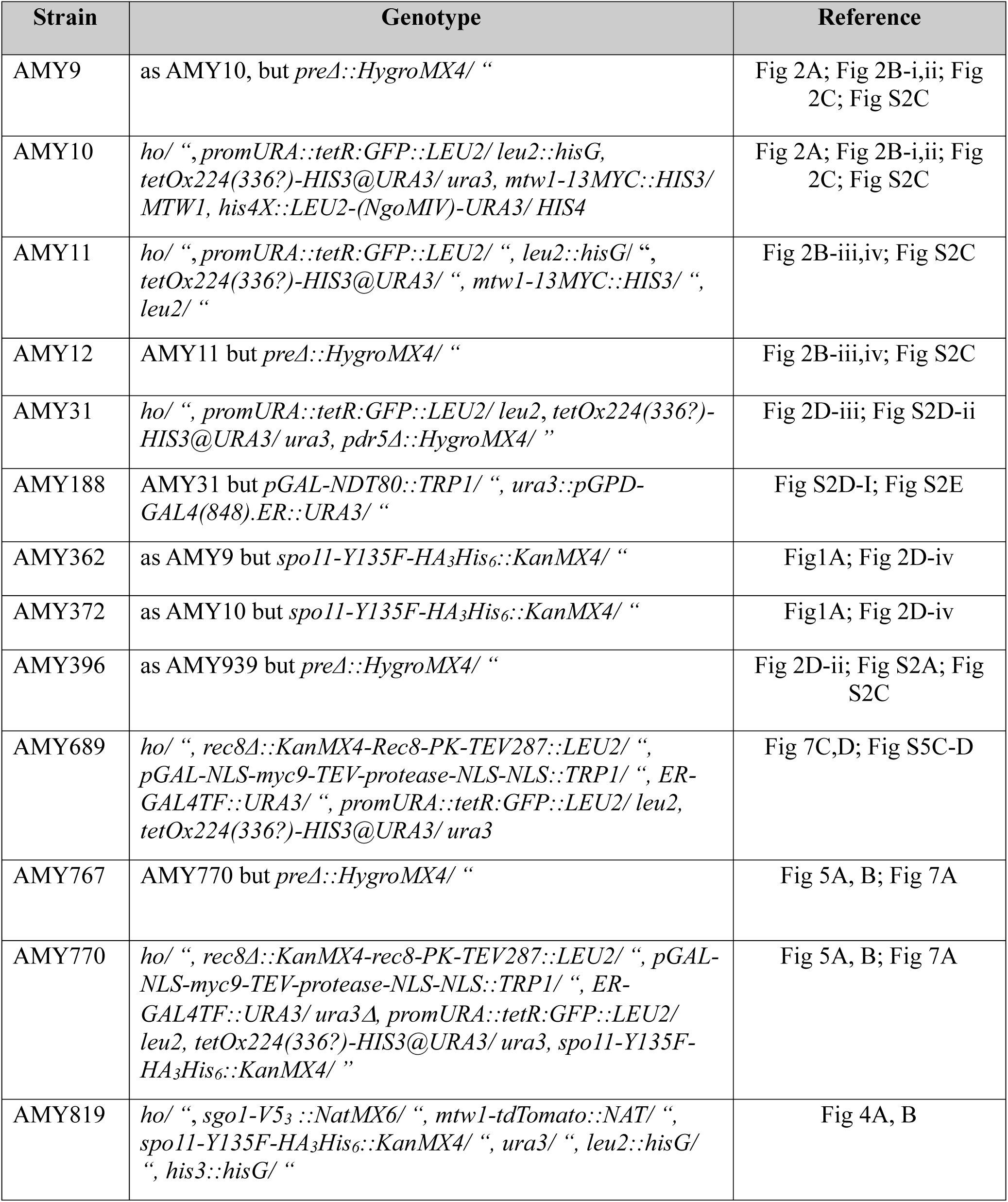
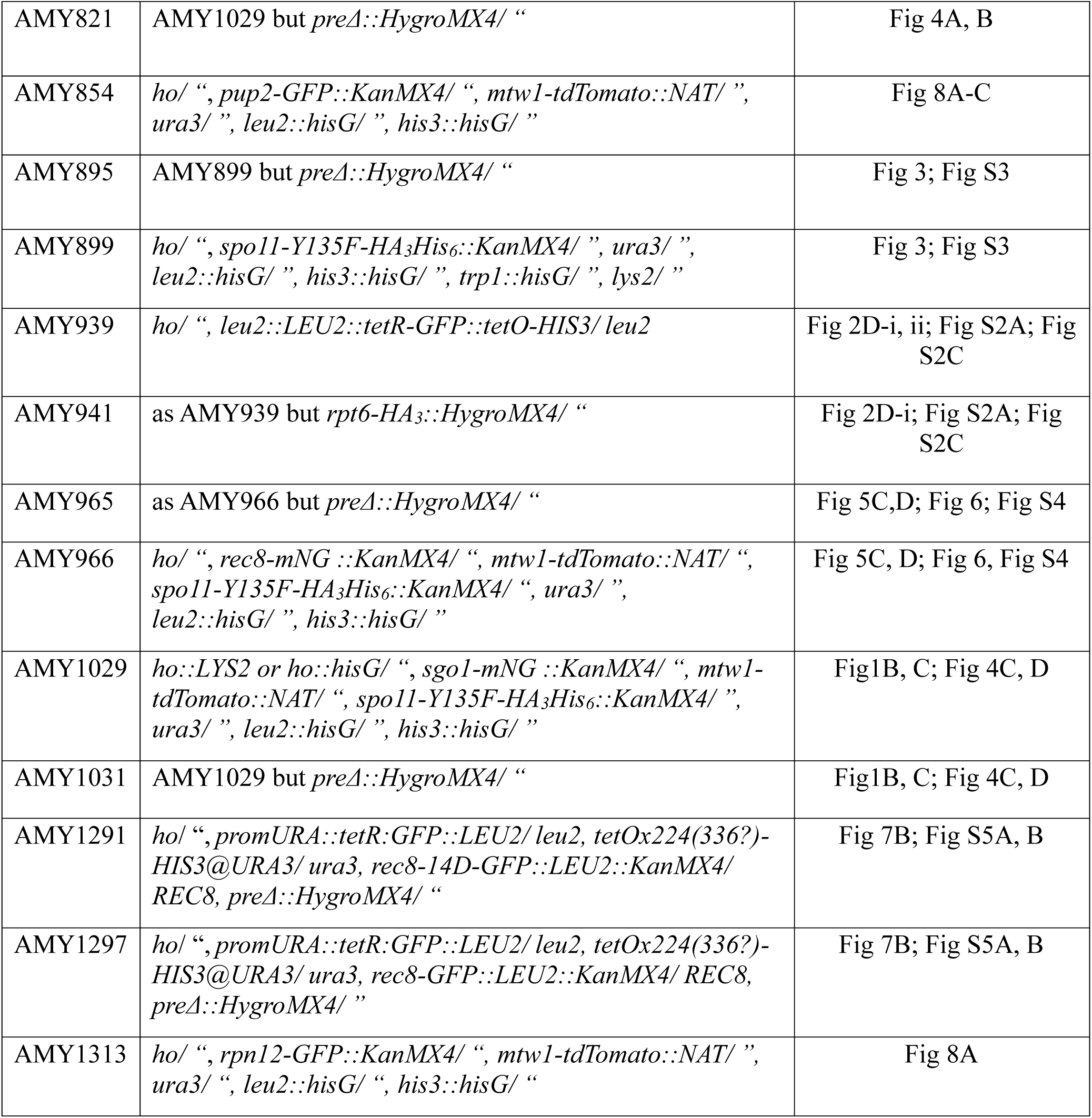

